# Cis-regulatory similarities in the zebrafish and human pancreas uncover potential disease-related enhancers

**DOI:** 10.1101/2020.04.27.064220

**Authors:** R. Bordeira-Carriço, J. Teixeira, M. Duque, M. Galhardo, D. Ribeiro, R. Dominguez-Acemel, P. N. Firbas, J. J. Tena, A. Eufrasio, J. Marques, F. Ferreira, T. Freitas, F. Carneiro, J. L. Goméz-Skarmeta, J. Bessa

## Abstract

The pancreas is a central organ for human diseases that have a dramatic societal burden, such as pancreatic cancer and diabetes^1,2^. Non-coding cis-regulatory elements (CREs) of DNA control gene expression^3,4^, being required for proper pancreas function. Most disease-associated alleles^5,6^ are non-coding, often overlapping with CREs^5^, suggesting that alterations in these regulatory sequences contribute to human pancreatic diseases by impairing gene expression. However, functional testing of CREs *in vivo* is not fully explored. Here we analysed histone modifications, transcription, chromatin accessibility and interactions, to identify zebrafish pancreas CREs and their human functional equivalents, uncovering disease-associated sequences across species. We found a human pancreatic enhancer whose deletion impairs the tumour suppressor gene *ARID1A* expression, conferring a potential tumour suppressor role to this non-coding sequence. Additionally, we identified a zebrafish *ptf1a* distal enhancer which deletion generates pancreatic agenesis, demonstrating the causality of this condition in humans^7^ and the interspecies functional equivalency of enhancers.

## Results

The pancreas of zebrafish, a vertebrate model suitable for genetic manipulation^8^, shares many similarities with the human pancreas, including transcription factors (TFs) operating in similar genetic networks of pancreas development and function^9,10^. We observed that these similarities are extensible to organ structure (Fig.1a) and cellular composition (SupplementaryFig.1), suggesting that shared genetic networks might operate through equivalent sets of CREs in both species.

**Figure 1.**
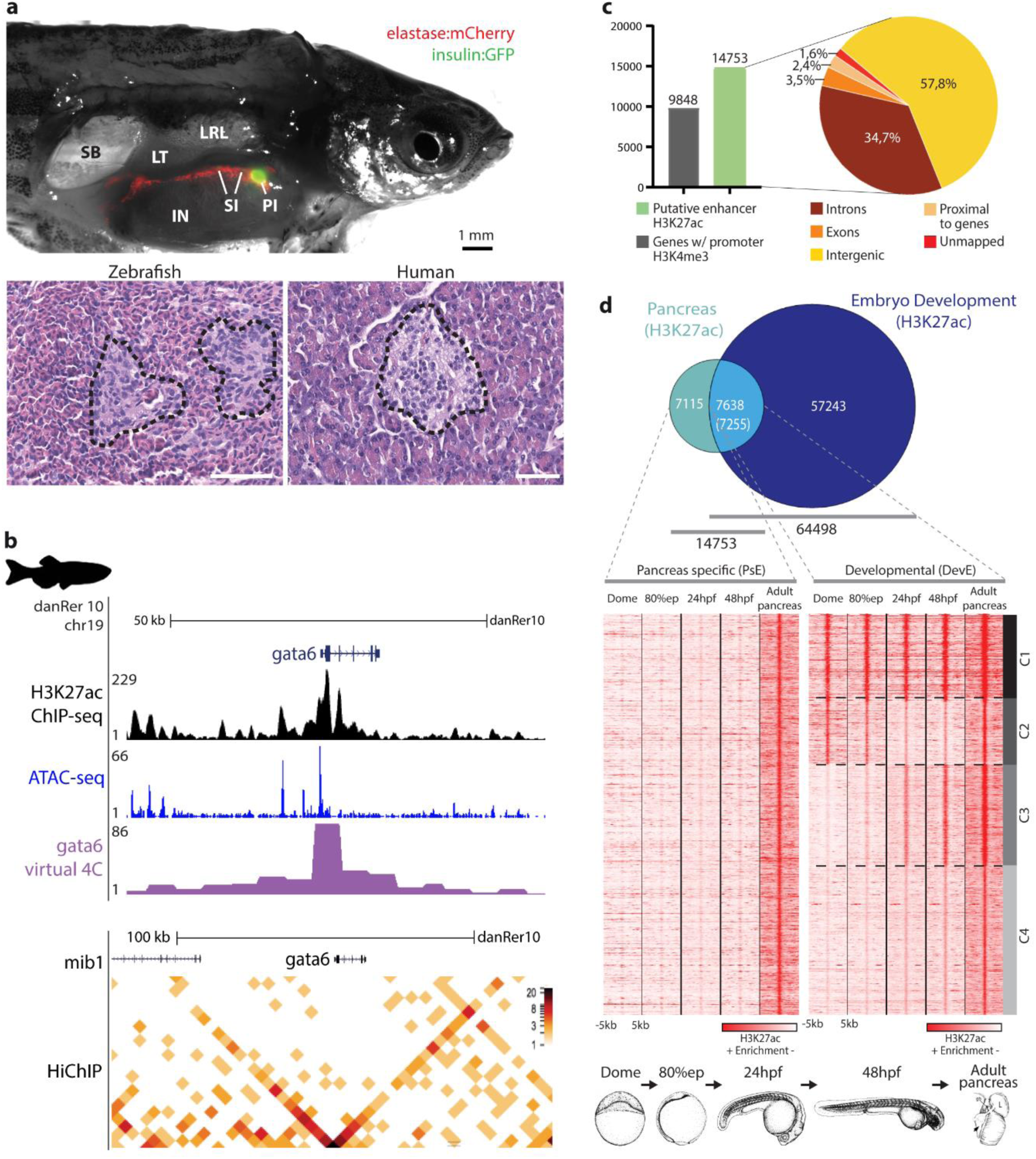
The zebrafish pancreas description, from histology to chromatin state. **a)** Comparison of adult zebrafish and human pancreas. Dissected adult male Tg(*insulin:GFP, elastase:mCherry*) zebrafish (top); insulin and elastase promoters drive GFP expression in beta-cells (green) and mCherry in acinar cells (red), respectively. IN, intestine; LRL, Liver right lobe; LT, left testis; PI, principal islet; SI, secondary islets; SB, swim bladder. Histology of the pancreas, transverse sections with hematoxylin/eosin staining (bottom). Islets of Langerhans (black dashed lines) surrounded by exocrine tissue in zebrafish (left) and human (right), magnification 40x. **b)** Genomic landscape of *gata6* in zebrafish pancreas with H3K27ac ChIP-seq profile (black), ATAC-seq peaks (blue) and *gata6* virtual 4C (purple) based on the interactions with *gata6* promoter detected by HiChIP for H3K4me3 (lower panel). **c)** Barplot (left) showing the number of genes with active promoters defined with H3K4me3 signal (gray bar) and putative active enhancers in adult zebrafish pancreas defined by H3K27ac mark (green bar), and their distribution throughout the regions of the genome (right). **d)** Venn diagram showing the overlap of putative enhancers in adult zebrafish pancreas and zebrafish embryonic development stages (top). Heat maps showing clusters of H3K27ac mark in dome, 80% epiboly (80%epi), 24hpf, 48hpf and adult pancreas for pancreas-specific enhancers (PsE) and developmental shared enhancers (DevE). A window of 10 kb around the reference coordinates for each sequence was used and the density files were subjected to k-means clustering, obtaining four different clusters in DevE: C1 - Cluster 1, C2 - Cluster 2, C3 - Cluster 3 and C4 - Cluster 4.

To identify CREs active in the zebrafish adult pancreas, we performed ChIP-seq for H3K27ac^11^, a key histone modification associated to active enhancers and ATAC-seq^12^, to identify regions of open chromatin. We have also performed HiChIP^13^ against H3K4me3^14^ to determine active promoters interacting with the uncovered enhancers (Fig.1b). We found 14753 putative active enhancers, mostly in intergenic regions (57.8%), and 23298 putative active promoters corresponding to 9848 genes (Fig.1c; SupplementaryTable1-3). To identify a subset of pancreas enhancers with higher tissue-specificity, we asked which of them are inactive in whole embryos (dome to 48hpf), finding that 7115 (48.2%) are active only in the differentiated adult pancreas (PsE; Fig.1d; SupplementaryTable4) while the remaining 7638 (51.8%) are also active during embryonic development (DevE). Interestingly, DevE presented 4 clusters (C1-4) with different activity dynamics during development (Fig.1d; SupplementaryFig.2; SupplementaryTable5).

Pancreatic enhancers should activate the expression of genes in the pancreas. To test this, we identified the nearest genes to each putative pancreas enhancer^15^, observing that genes nearby PsE are enriched for exocrine pancreas expression (p<4.27e-9; SupplementaryFig.3a; SupplementaryTable6-7), detected by *in situ* hybridization. These results contrast with DevE, further suggesting a higher tissue-specificity of PsE. To improve the enhancer to gene association, we used H3K4me3 HiChIP to detect interactions between active promoters and putative enhancers in the zebrafish adult pancreas (Fig.1b; SupplementaryTable8), and used RNA-seq to evaluate transcription. We found that PsE-associated genes have a higher average expression in a variety of pancreatic cell types when comparing to all transcribed genes, contrasting with transcription in the muscle (Fig.2a, SupplementaryTable9). Similar results were obtained when analysing genes associated to the remaining clusters of pancreatic enhancers (PsEs+DevE, DevE and C1-4; SupplementaryFig.3). Performing a similar assay using the transcriptome of whole zebrafish embryos from 18 developmental stages^16^, genes associated to DevE have shown an increased average expression comparing to all transcribed genes, with a similar dynamic to the enhancer activation during development (Fig.2b; SupplementaryFig.4). These results suggest that DevE enhancers control gene expression in the adult differentiated pancreas and during development.

**Figure 2.**
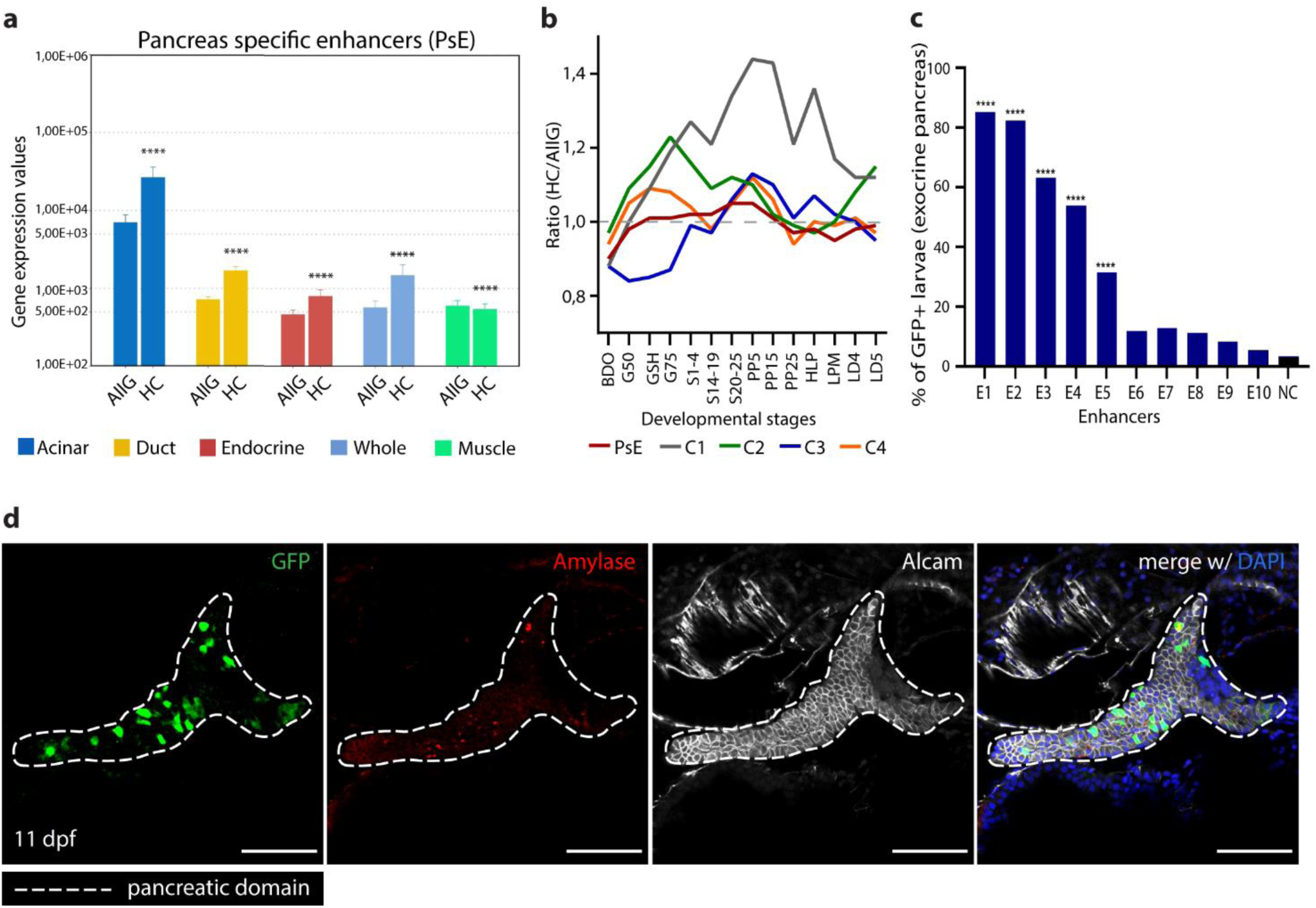
ChIP-seq and ATAC-seq data accurately predict functional pancreatic enhancers. **a)** Average expression of genes interacting with pancreas-specific enhancers detected by HiChIP for H3K4me3 (HC), compared to the average expression of all genes (AllG). Gene expression was determined from RNA-seq data from different pancreatic cells (acinar, duct ^62^, endocrine and whole pancreas) and muscle (control). Wilcoxon test, *p*-values ≤ 0.05 were considered significant. \*\*\*\**p*<0,0001. **b)** Ratio between the average expression of genes interacting with pancreas-specific enhancers (PsE, C1, C2, C3 and C4 clusters) and the average expression of all genes throughout zebrafish development. BDO: blastula, dome; G50: gastrula, 50%epiboly; GSH: gastrula,shield; G75: gastrula,75%epiboly; S1-4: segmentation, 1-4somites; S14-19: segmentation, 14-19somites; S20-25: segmentation, 20-25somites; PP5: pharyngula, prim5; PP15: pharyngula, prim15; PP25: pharyngula, prim25; HLP: hatching, long pec; LPM: larval, protruding mouth; LD4: larval, day4; LD5: larval, day5. **c)** Percentage of zebrafish larvae showing GFP expression in the pancreas after *in vivo* transient transgenesis reporter assays with Z48 vector. A total of 10 putative enhancer sequences (E1 to 10) were tested, using the empty enhancer reporter vector as negative control (NC). Values are represented as percentages and compared by Chi-square test. p-values<0.05 were considered significant (****p<0.0001). **d)** Representative confocal image of the *in vivo* transient transgenesis reporter assays for the E3 sequence (zPtf1aE1) showing expression of GFP (green) in 11dpf zebrafish pancreas (white dashed line), labelled by anti-Alcam staining (white) and anti-Amylase (red) antibodies. Nuclei were stained with DAPI (blue). Images were captured with a Leica SP5II confocal microscope. Scale bar 60 µm.

To determine if the identified regulatory sequences are active pancreatic enhancers, we have performed *in vivo* reporter assays for 10 regions with strong H3K27ac signal and 7 with low levels of this mark (Fig.2c-d; Supplementary Fig.5). From the first set, we have found that 5 out of 10 tested sequences (H3K27ac: −log10(*p*-value)≥35) are pancreatic enhancers (50%; Fig.2c-d and SupplementaryTable10). In contrast, from the regions with low H3K27ac signal, only 1 out of 7 tested (14%, Supplementary Fig.5) showed strong and reproducible pancreatic enhancer activity (−log10(*p*-value)<35). These results validate the robustness of the enhancers prediction based on chromatin state.

We observed that out of 14753 putative zebrafish pancreas enhancers, only 12.49% (n=1842) could be directly aligned to the human genome^17^ (Fig.3a; SupplementaryTable11), a similar proportion found in developmental enhancers (11.36%; 7326 out of 64498), with both groups sharing similar PhastCons conservation scores (Fig.3b; SupplementaryFig.6a; SupplementaryTable11). Next, we wanted to discern if the zebrafish putative pancreas enhancers that align to the human genome also overlap with H3K27ac signal from human pancreas. Although only a minority of interspecies aligned sequences shared H3K27ac signal (Total pancreas data set: 229 out of 1842; PsE: 116 out of 1052; DevE: 113 out of 790), there is a clear enrichment comparing to random sequences (Fig.3c), although not showing a higher average sequence conservation score (Fig.3b). These results suggest that pancreatic enhancer function is not a strong constraint for sequence conservation.

**Figure 3.**
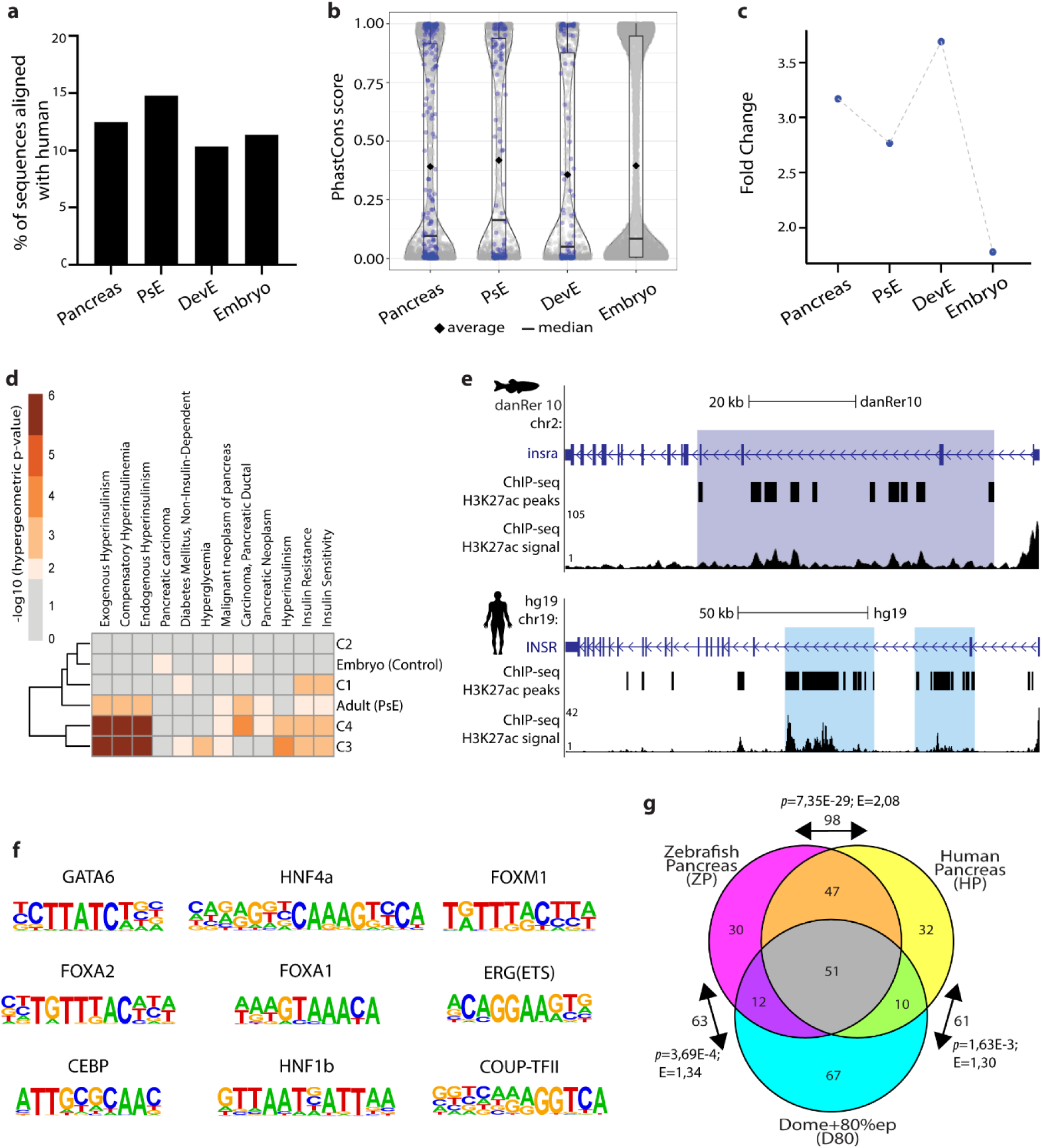
The zebrafish and human pancreas share cis-regulatory similarities. **a)** Percentage of predicted zebrafish pancreatic enhancer sequences that can be aligned to the human genome. Sequences are grouped in different clusters: putative enhancer sequences active in total pancreas (Pancreas), active only in differentiated adult pancreas (PsE), active in differentiated adult pancreas and during embryonic development (DevE) and active only during embryonic development (Embryo). **b)** PhastCon scores (99 vertebrate genomes) for human sequences aligned to zebrafish putative enhancers (Pancreas, PsE, DevE and Embryo; grey). In blue are depicted sequences that also show H3K27ac signal in human pancreas (data from ENCODE). **c)** Fold change of the number of zebrafish pancreatic H3K27ac positive sequences that overlap with human pancreatic H3K27ac positive sequences (ENCODE data), comparing to a 10^5 random shuffle of the used human sequences. **d)** Heat map showing −log10(p-values) from hypergeometric tests of the enrichment for pancreatic disease association on the genes linked to each enhancer cluster identified by HiChIP for H3K4me3. The coloured values meet the established criteria of statistical significance (q-value≤0.05) and fold change (abs(FC)≥1.5). **e)** Genomic landscape of the human *INSR* gene (top) and zebrafish *arid1ab* ortholog (bottom), showing H3K27ac signal and the position of predicted super-enhancers highlighted in blue. **f)** List of known relevant pancreas transcription factors (TFs) whose motifs are enriched in zebrafish pancreas H3K27ac ChIP-seq data. **g)** Venn diagram of the top 140 enriched TFBS motifs detected in H3K27ac positive sequences in three different datasets: zebrafish pancreas (ZP), human pancreas (HP) and dome+80%epiboly embryos (D80). Arrows show the number of motifs shared between pairs of groups. Statistical significance was determined by hypergeometric enrichment test, *p*-values and the enrichment of the observed *vs* expected are represented.

Then, we wanted to assess whether functionally equivalent pancreatic CREs might exist between human and zebrafish, despite an overall lack of sequence conservation. To explore this possibility, we analysed if the human ortholog genes coupled to each cluster of zebrafish enhancers were enriched for human pancreatic diseases. Such enrichment was observed for the clusters of late development and adult (PsE, C3 and C4; Fig.3d; SupplementaryTable12). Subsequently, we searched for super enhancers active in the pancreas of both species, finding 275 in zebrafish and 875 in humans (SupplementaryTable13). Gene ontology for putative target genes showed a similar enrichment for transcriptional regulation in both species and several of these genes corresponded to the same orthologs (32 out of 271-zebrafish, SupplementaryFig.6b-c), some having important pancreatic functions, such as *INSR* and *GATA6* (Fig.3e; SupplementaryFig.6d). We further asked if human and zebrafish enhancers might operate similarly, using equivalent TFs. To test this, we performed motif discovery for TF binding sites (TFBS) in regions of open chromatin identified by ATAC-seq^12^, within the 14753 pancreas enhancers, finding several TFBS for known pancreatic TFs (ZP; Fig.3f; SupplementaryTable14). We have performed a similar analysis using available human whole pancreas datasets (HP) and, to establish comparisons, for zebrafish embryos (D80, dome and 80%epiboly; 24HPF, 24 hpf) and human heart ventricle (V). Selecting the top 140 enriched motifs from each dataset, we observed that the majority of the common motifs were found in zebrafish (ZP) and human (HP) pancreas datasets (ZP,HP:98; ZP,D80:63; HP,D80:61), while comparisons with the human ventricle (V) showed that ZP,HP was the second largest group (Fig.3g; SupplementaryFig.7a-b). Additionally, selecting 25 motifs from TFs known to be required for pancreas function or development, we found that the majority of those were within the ZP,HP overlapping datasets, regardless of the compared groups (SupplementaryFig.7c-e). These results suggest that the same set of TFs might operate in zebrafish and human pancreas enhancers. Overall, these results argue in favour of interspecies functional equivalency of enhancers. To better address this hypothesis, we focused on the regulatory landscape of *arid1ab*, the orthologue of human *ARID1A*, a tumour-suppressor gene known to be associated with cancer in several organs, including pancreas^18,19^. We identified one robust zebrafish pancreatic enhancer (zArid1abE3, Fig.4a), validated *in vivo* (Fig.4b; SupplementaryFig.8), that interacts with the promoter of *arid1ab* (Fig.4a; SupplementaryFig.9). Additionally, we detected a human/zebrafish syntenic block containing the zebrafish zArid1abE3 enhancer and a human pancreatic CRE (hArid1abE3). Enhancer assays for hArid1abE3 demonstrated its ability to drive expression in the pancreas, suggesting a functional equivalency to the zebrafish zArid1abE3 enhancer (Fig.4b; SupplementaryFig.8). To study the impact of this human enhancer in ARID1A expression, we deleted hArid1abE3 enhancer in hTERT-HPNE cell line, a human pancreatic duct cell line, through CRISPR-cas9 system (SupplementaryFig.10), using as a control a deletion in an unrelated genomic region^20^. We observed lower ARID1A expression upon deletion of hAridE3 than in the control (Fig4c-e and SupplementaryFig10), suggesting that the loss of this enhancer might interfere with the DNA-damage response, with possible implications in the increased risk for pancreatic cancer^21,22^.

**Figure 4.**
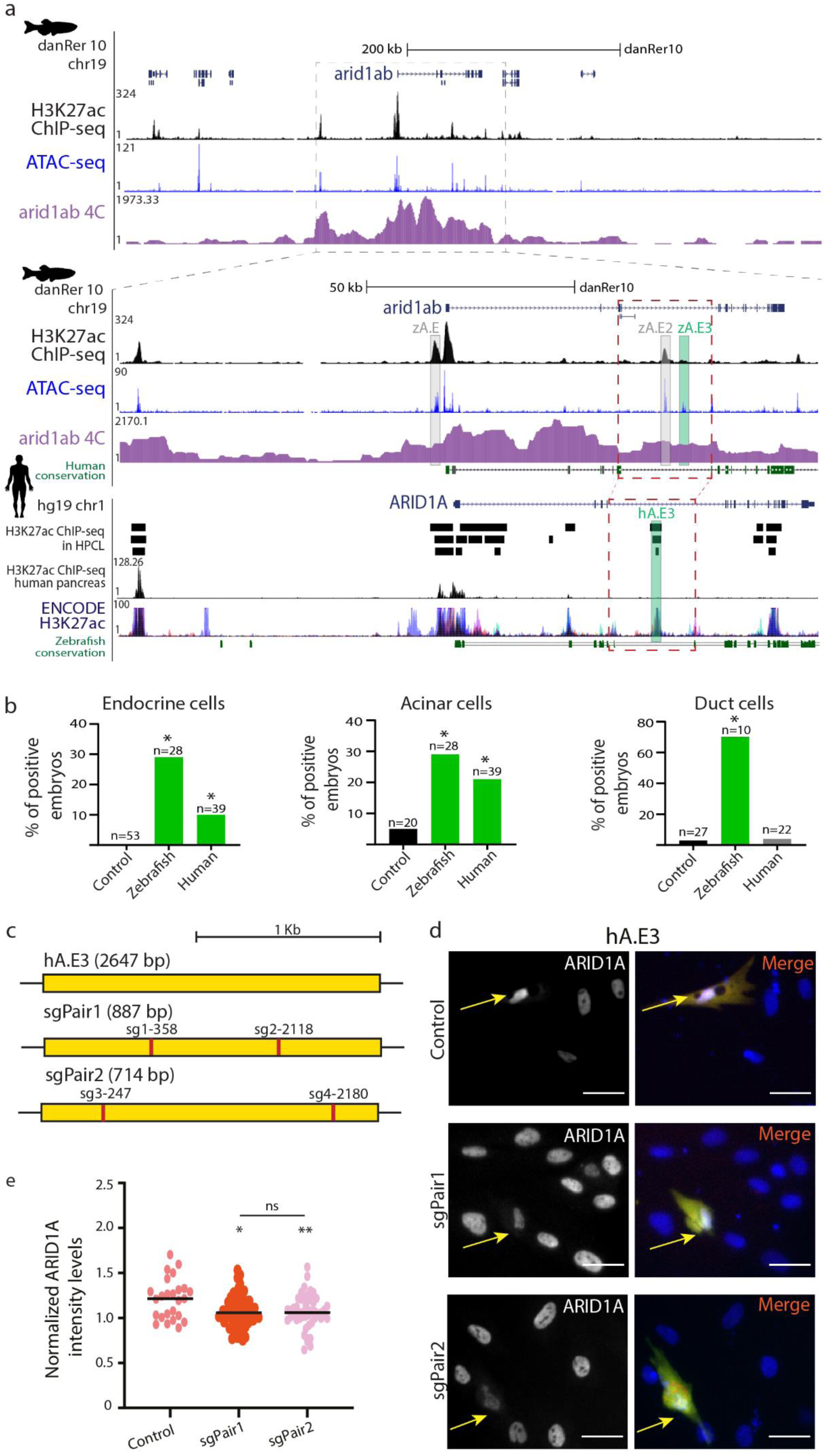
The zebrafish and human *arid1ab/ARID1A* regulatory landscapes contain an equivalent pancreatic enhancer. **a)** Genomic landscape of the zebrafish *arid1ab* gene, showing profiles for H3K27ac ChIP-seq (black), ATAC-seq (blue) and 4C with view point in the *arid1ab* promoter (purple) in adult zebrafish pancreas (upper panel); zoom-in of the *arid1ab* gene and putative enhancers in its vicinity (middle panel). Genomic landscape of human *ARID1A* (lower panel), with annotated H3K27ac enriched intervals based on peak calling from different human pancreatic cell lines (HPCL, black bars), PT-45-P1, CFPAC-1 and HPAF-II, top to bottom, respectively, H3K27ac profile from human whole pancreatic tissue (WPT, black) and H3K27ac profile from non-pancreatic human cell lines (NPHCL; GM12878, H1-hESC, HSMM, HUVEC, K562, NHEK and NHLF; Data from ENCODE). Sequence conservation between human and zebrafish is represented in dark green. Tested putative enhancers from zebrafish and human genomic DNA are highlighted in grey zA.E1 (zAridE1) and zA.E2 (zAridE2) and green zA.E3 (zAridE3) and hA.E1 (hAridE3). A zebrafish/human syntenic box containing the zAridE3 sequence and its putative equivalent human enhancer hAridE3 is highlighted (red dashed box). **b)** Percentage of zebrafish embryos showing GFP zAridE3 or hAridE3 mediated expression from transient transgenesis assays in endocrine, acinar and duct cells, at 11dpf. Statistical significance determined by Chi-square test with Yates correction (*p<0.05). **c)** Schematic representation of the targeting strategy for the hAridE3 locus, indicating the CRISPR sgRNA target sites and expected genomic deletions. **d)** Representative images of transfected hTERT-HPNE human cells assays. In the control, a region without H3K27ac ChIP-seq signal was deleted. In sgPair1 and sgPair2, the hAridE3 (hA.E3) locus was deleted using the respective pairs of sgRNAs. The double transfected cells are defined by GFP (green) and mCherry (red) co-expression and the nuclei were stained with DAPI (blue) and anti-ARID1A (grey). The yellow arrow is pointing to the double transfected cells. Images were captured with Leica DMI6000 FFW microscope. Scale bar: 40 μm. **e)** Normalized ARID1A intensity levels, measured from immunocytochemistry images, in control cells and in sgPair1 and sgPair2, where hA.E3 locus was deleted. Statistical significance is depicted in the graphs for p≤0.05, p≤0.01(**) and ns for no statistical significance. These results were obtained from three batches of independent experiments.

To further evaluate the interspecies functional equivalency of enhancers, we have focused in the human locus of *PTF1A*, known to be controlled by a distal downstream enhancer whose deletion was associated to pancreatic agenesis^7^ (Fig.5a; hPtf1aE3). Concomitantly, we detected a novel putative zebrafish enhancer located distally downstream of *ptf1a* (zPtf1aE3), apart from two previously identified proximal enhancers (zPtf1aE1 and zPtf1aE2)^23^. zPtf1aE3 interacts with the promoter of *ptf1a*, observed by Hi-ChIP and 4C-seq (Fig.5a; SupplementaryFig.9), and could correspond to the functional equivalent enhancer associated with pancreatic agenesis in humans (hPtf1aE3). *In vivo* enhancer assays for zPtf1aE3 and hPtf1aE3 have shown that both are pancreatic enhancers able to drive expression in the same cell types (acinar, duct and pancreas progenitor cells), with a more robust expression in progenitor cells (Fig.5b; SupplementaryFig.11-12), suggesting that these two enhancers share some regulatory information. This is further supported by binding sites for FOXA2 and PDX1 in the human hPtf1aE3, also predicted to bind to the zebrafish zptf1aE3 (SupplementaryFig.13). To further validate these results, we have generated genomic deletions in the zPtf1aE3 sequence, observing a reduction in the pancreatic progenitor domain, resulting in pancreatic hypoplasia (Fig.5d; SupplementaryFig.14-16), compatible with loss-of-function of *ptf1a* in zebrafish^23^ and the proposed loss of hPtf1aE3 function in humans^7^. These results demonstrate that zebrafish and humans share a functional equivalent distal enhancer of *PTF1A*, clarifying the causality of pancreatic agenesis upon enhancer disruption in humans.

**Figure 5.**
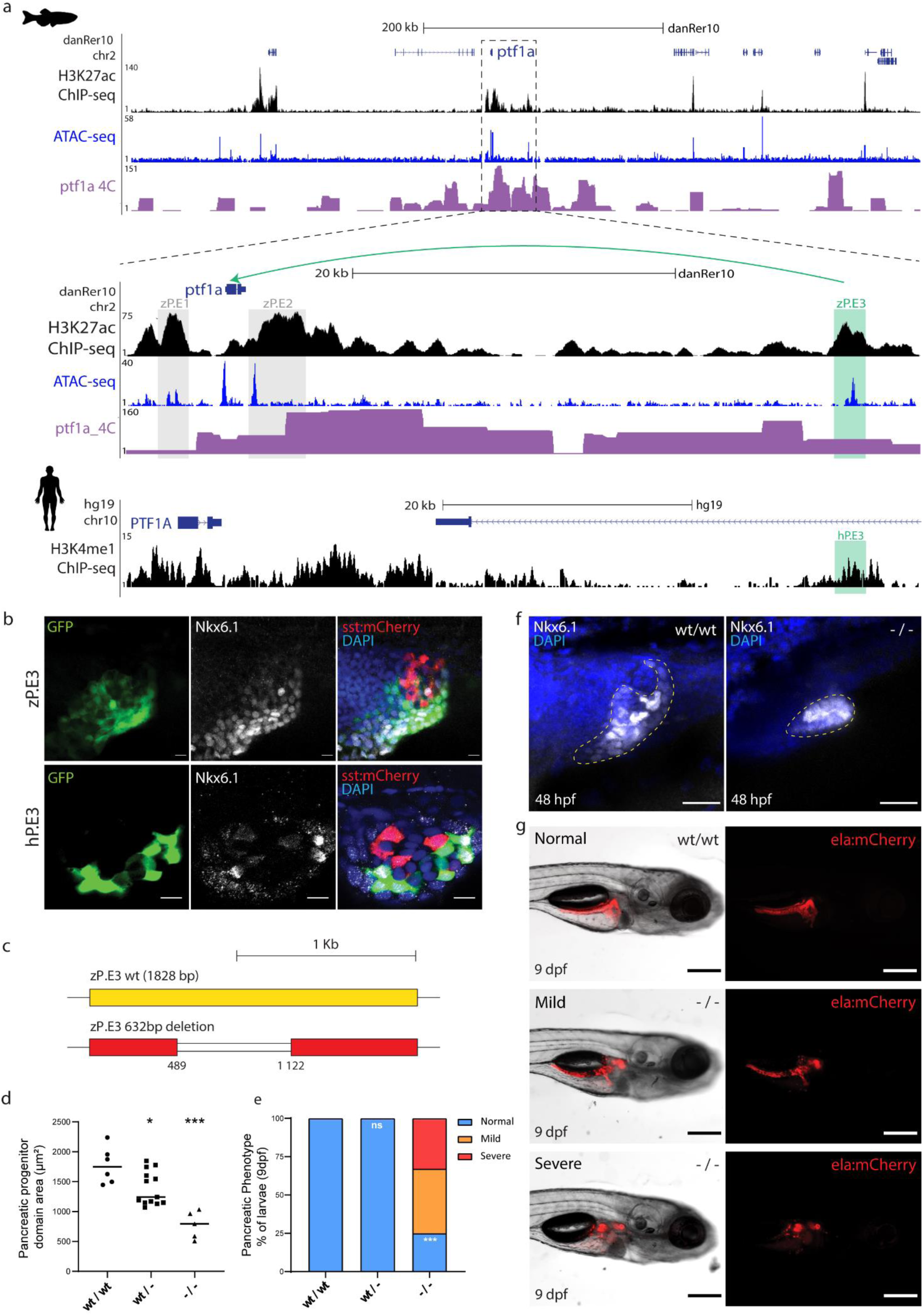
The *Ptf1a* regulatory landscape in zebrafish and human contains a functional equivalent enhancer essential for pancreas formation. **a)** UCSC Genome Browser view of the zebrafish *ptf1a* and human *PTF1A* genomic landscapes showing H3K27ac ChIP-seq (black), ATAC-seq (blue) and ptf1a 4C interactions (purple) from whole zebrafish pancreas samples (upper panel), with a zoom-in (middle panel), and H3K4me1 ChIP-seq data (black) from human embryonic pancreatic progenitors (lower panel). Grey boxes highlight two previously validated zebrafish enhancers, zP.E1 (zptf1aE1) and zP.E2 (zptf1aE2) in the vicinity of the *ptf1a* gene. Green boxes highlight a newly discovered distal enhancer in zebrafish (zP.E3 (zptf1aE3), and the location of its putative human functional ortholog (hP.E3). **b)** Confocal images of zebrafish reporter transgenic lines Tg(zP.E3:GFP) and Tg(hP.E3:GFP), showing co-localization of GFP expression (green) with anti-NKX6.1 (white), a marker of pancreatic progenitors, at 48 hpf. Delta-cells of the endocrine pancreas express mCherry (red) and nuclei are labelled with DAPI (blue). Scale bar 25 μm. **c)** Schematic depiction of the CRISPR/Cas9 mediated 632 bp deletion of the zP.E3 enhancer. **d)** Pancreatic progenitor domain area, defined by anti-NKX6.1 (white), of homozygous (-/-), heterozygous (wt/-) and wild type (wt/wt) embryos for the deletion of zP.E, at 48 hpf, (n=24). **e)** Percentage of larvae, -/-, wt/- and wt/wt, with different pancreatic phenotypic defects (normal, mild and severe) at 9 dpf, (n=47). Fisher’s exact test, p-values<0.05 were considered significant (***p<0.001, *p<0.05). **f)** Representative confocal images (maximum intensity projections) of the pancreatic progenitor domain (NKX6.1, white) of zP.E3wt/wt and zP.E3-/- embryos at 48 hpf. Nuclei are stained with DAPI. Scale bar: 25 μm. **g)** Epifluorescence live images of representative phenotypes quantified in e), for Tg(elastase:mCherry) zebrafish larvae at 9dpf. Scale bar: 250 μm.

Cis-regulatory mutations and variations are associated with pancreatic cancer and diabetes ^5,24–27^, however the *in vivo* implications of these genetic changes are yet unknown. Here we explore the chromatin state of the zebrafish pancreas, to uncover pancreatic enhancers and establish comparisons with humans, to predict and model human pancreas disease-associated enhancers. We found that, although most of the zebrafish pancreas enhancers show no sequence conservation with human pancreas enhancers, they share many TFBS and their target genes are enriched for human pancreas diseases. These results suggest the existence of functionally equivalent enhancers in zebrafish and humans, as proposed for other tissues and species^28^. We illustrate this finding for the regulatory landscape of *ARID1A*, a tumour-suppressor gene active in the pancreas^21,29^ and other tissues^18^, finding a human pancreatic enhancer, whose deletion impairs *ARID1A* expression, defining a locus for non-coding mutations that might increase the risk for pancreatic cancer. We further validated these findings for one enhancer of *PTF1A*^30^, where both zebrafish and human enhancers share regulatory information and biological requirement, showing a decrease in the pancreatic progenitor domain in its absence and, as a consequence, developing pancreatic agenesis. In summary, we show that transcriptional cis-regulation of the human and zebrafish adult pancreas have a high similarity, allowing the functional exploration of cis-regulatory sequences in zebrafish, with the potential of translation to human pancreatic diseases.

## Acknowledgements

This study was supported by the *European Research Council (ERC) under the European Union’s Horizon 2020 research and innovation programme* (grant agreement no. ERC-2015-StG-680156-ZPR and ERC-2016-AdG-740041-EvoLand to JLG-S). JB acknowledges Fundação para a Ciência e a Tecnologia (FCT), for a FCT Investigator position (Grant IF/00654/2013). JLG-S also acknowledges the Spanish Ministerio de Economía y Competitividad (Grant BFU2016-74961-P), the Marató TV3 Fundacion (Grant 201611) and the institutional grant Unidad de Excelencia María de Maeztu (MDM-2016-0687 to the Department of Gene regulation and morphogenesis of Centro Andaluz de Biología del Desarrollo. RBC was funded by FCT (ON2201403-CTO-BPD), IBMC (BIM/04293-UID991520-BPD) and EMBO (Short-Term Fellowship). JTx (SFRH/BD/126467/2016), MD (SFRH/BD/135957/2018), AE (SFRH/BD/147762/2019), and FF (PD/BD/105745/2014) are PhD fellows from FCT. MG was supported by the EnvMetaGen project via the European Union’s Horizon 2020 research and innovation programme (grant agreement 668981). Authors acknowledge the support of i3S Scientific Platform Advanced Light Microscopy, member of the national infrastructure PPBI-Portuguese Platform of BioImaging (supported by POCI-01-0145-FEDER-022122). Authors acknowledge Carla Oliveira (Microenvironments for New Therapies) for statistical support, Catarina Meireles and Emilia Cardoso from the Translational Cytometry (TraCy) Scientific Platform, Mafalda Sousa for the automated cell analysis, Paula Magalhães and Tânia Meireles from Cell Culture and Genotyping Scientific Platform, Guilherme Cardoso from the Histology Service from Ipatimup Diagnostics, the I3s hpc facility used for data processing, in particular to Andre Torres, and finally Isabel Guedes for support in the maintenance of zebrafish lines. We also acknowledge, from CABD, the support from Elisa de la Calle-Mustienes and Sandra Jimenez Gancedo on ChiP-seq, Ana Fernandez-Minñán on 4C-seq and Ensieh Farahani on ATAC-seq protocols.

## Author Contributions

JB designed the study with input of RBC and JLGS. JB coordinated the project. JB and JLGS supervised the work. All authors contributed for the development and discussion of the work. RBC obtained biological material and generated next-generation sequencing data from zebrafish pancreas. JT, MG, RDA, PNF and JJT performed computational analyses and data interpretation. RBC, MD, AE and DR performed enhancer-assays in zebrafish, from plasmid generation to transgenesis, and Crispr-Cas9 in zebrafish. MD, DR and AE performed immunohistochemistry, microscopy acquisition and analysis. JTx performed transfection, Crispr-Cas9 and image acquisition in human cell lines with support of FF. FC contributed with histology of human pancreas. TF and JM contributed for plasmid and zebrafish lines generation. JB wrote the manuscript with input from all authors.

## Material and Methods

### I) Experimental procedures

#### Zebrafish stocks, husbandry, breeding and embryo culture

Adult zebrafish AB/TU WT strains, transgenic and mutant lines were maintained at 26-28°C under a 10h dark/14h light cycle in a recirculating housing system according to standard protocols^31^. Embryos were grown at 28°C in E3 medium [5mM NaCl, 0.17mM KCl, 0.33mM CaCl_2_•2H_2_O, 0.33mM MgSO_4_•7H_2_O and 0.01% methylene blue (Sigma-Aldrich), pH 7.2] or E3 supplemented with 0.01% PTU (1-phenyl-2-thiourea)^32^. For the *in vivo* enhancer assays, embryos were anesthetized by adding tricaine (MS222; ethyl-3-aminobenzoate methanesulfonate, #E10521-10G, Sigma-Aldrich) to the medium and selected by the internal positive control of transgenesis. For the establishment of transgenic and mutant zebrafish lines, embryos were microinjected, selected, bleached and grown until adulthood. Adult F0s were outcrossed with WT adults and the offspring screened for the internal control of transgenesis and the pattern of expression of the regulatory element, or for the respective mutations, by genotyping. *In vivo* reporter lines, Tg(ela:mCherry) and Tg(sst:mCherry), were used to label the exocrine and endocrine domain, respectively. The i3S animal facility and this project were licensed by *Direcção Geral de Alimentação e Veterinária (DGAV)* and all the protocols used for the experiments were approved by the i3S Animal Welfare and Ethics Review Body.

#### Cell culture

hTERT-HPNE (intermediary cells formed during acinar-to-ductal metaplasia; ATCC CRL-4023) cells were cultured in a 5% CO_2_-humidified chamber at 37°C in DMEM (1x, 4.5 g/L D-glucose with pyruvate; #D6429, Gibco, ThermoFisher Scientific), supplemented with 10% fetal bovine serum (#BCS0615, biotecnomica), 10ng/mL human recombinant EGF (#11343406, Immunotools) and 750ng/mL puromycin (#P8833-25MG, Sigma-Aldrich) in TC Dish 100 (SARSTEDT). When cells reached to 90% of confluence, they were split using TrypLE Express (#12604-021, Gibco, ThermoFisher Scientific; approximately 0.5 mL per 10 cm2).

#### ChIP-seq

Whole pancreas was dissected from 25 adult zebrafish (∼50×10^6^ cells), kept on ice in PBS with 1x Complete Proteinase Inhibitor (#11697498001, Roche), fixed in 2% formaldehyde (#F1635-500ML, Sigma-Aldrich) for 10 min, and stored at −80°C. ChIP was performed as previously described for zebrafish embryos^33^ with minor alterations. Cell lysis was performed on ice, using a 15 mL Tenbroeck Homogenizer, in cell lysis buffer [10mM Tris-HCl pH7.5, 10mM NaCl, 0.5% NP-40, 1x Complete Proteinase Inhibitor (#11697498001, Roche)] for 15 min. Nuclei were washed and re-suspended in nuclei lysis buffer (50mM Tris-HCl pH7.5, 10mM EDTA, 1% SDS, 1x Complete Proteinase Inhibitor (#11697498001, Roche)). Chromatin was sheared using a BioruptorPlus (Diagenode) device with the following cycling conditions: 10 min high–30 sec on, 30 sec off; 15 min on ice; 10 min high–30 sec on, 30 sec off. The sonicated chromatin had a size in the range of 100–500 bp and was incubated overnight at 4°C with the anti-H3K27ac antibody (#ab4729, Abcam). Samples were incubated for 1h at 4°C with Dynabeads Protein G for Immunoprecipitation (#10003D, Invitrogen, ThermoFisher Scientific). Final DNA was purified with MinElute (#28004, Qiagen) and sequenced on Illumina HiSeq 2000 platform.

#### ATAC-seq

ATAC-seq was performed as previously described^34^, with minor changes. Whole pancreas was dissected from 2-3 adult zebrafish. Following cell lysis, 50000-10000 nuclei were submitted to tagmentation with Nextera DNA Library Preparation Kit (#FC-121-1030, Illumina). ATAC-seq libraries were amplified KAPA HiFi HotStart PCR Kit (Roche) with the primers Ad1, Ad2.2 and Ad2.3 ^12^, and further purified with PCR Cleanup Kit (#28104, Qiagen).

#### 4C-seq

4C-seq was performed as previously described^34^, with minor alterations. Whole pancreas was dissected from 6-12 adult zebrafish (7-15×10^6^ cells), kept on ice in PBS with 1x Complete Proteinase Inhibitor (#11697498001, Roche), fixed in 2% formaldehyde (#F1635-500ML, Sigma-Aldrich) for 10 min, and stored at −80°C. Cell lysis was performed on ice, with a 15 mL Tenbroeck Homogenizer, not exceeding 10 min. Ligation was performed with 60U T4 DNA Ligase (#EL0012, ThermoFisher Scientific). The restriction enzymes used were DpnII (#R0543M, NEB) and Csp6I (#ER0211, ThermoFisher Scientific) for the first and second cuts, respectively. Chromatin was purified by Amicon Ultra-15 Centrifugal Filter Device (Milipore). 4C libraries were prepared for Illumina sequencing by the Expand Long Template Polymerase (Roche) with primers targeting the TSSs of each gene and including Illumina adapters (SupplementaryTable16). Final PCR products were purified with the High Pure PCR Product Purification Kit (Roche) and AMPure XP PCR purification kit (Agencourt AMPure XP).

#### HiChIP-seq

HiChIP-seq was performed as previously described^13^, with minor alterations. Whole pancreas was dissected, fixed in 1% formaldehyde (#F1635-500ML, Sigma-Aldrich) and cells lysed as described for 4C-seq. Immediately after lysis, samples were washed with HiChIP Wash Buffer (Tris-HCl pH 8 50mM, NaCl 50 mM, EDTA 1 mM). Chromatin was sonicated using the BioruptorPlus (Diagenode) with the following cycling conditions: 10 min high–30 sec on, 30 sec off; 15 min on ice, to obtain a size in the range of 100–500 bp. Samples were incubated with anti-H3K4me3 antibody (#AB8580, Abcam) and Dynabeads Protein G for Immunoprecipitation (#10003D, Invitrogen, ThermoFisher Scientific) and purified with DNA Clean and Concentrator columns (Zymo Research). Up to 150 ng of the DNA was then biotinylated with Streptavidin C-1 beads (ThermoFisher Scientific). Tagmentation was performed using Nextera DNA Library Preparation Kit (#FC-121-1030, Illumina). Libraries were amplified using NEBNext® High-Fidelity 2X PCR Master Mix (#M0541S, NEB) with primers Ad1, Ad2.23 and Ad2.24^12^. The final product was purified with DNA Clean and Concentrator kit (Zymo Research).

#### Generation of plasmids for enhancer assays

Putative enhancer sequences were selected based on the overlap between H3K27Ac ChIP-seq and ATAC-seq signal in non-coding regions within the landscape of each pancreas-relevant gene. Sequences were PCR amplified from zebrafish genomic DNA using the primers in SupplementaryTable16 (Sigma-Aldrich), with the proof-reading iMax ™ II DNA polymerase (INtRON Biotechnology) following the manufacturer’s instructions for a standard 20 μl PCR reaction. PCR products were visualized by electrophoresis on an 1% agarose gel, the bands excised, purified with NZYGelpure kit (NZYTech) and cloned into the entry vector pCR®8/GW/TOPO (#250020 Invitrogen, ThermoFisher Scientific) according with manufacturer’s instructions. The vectors were then recombined into the destination vectors Z48^35^, for transient enhancer assays, and ZED^36,37^, for stable transgenic lines, using Gateway® LR Clonase® II Enzyme mix (Invitrogen, ThermoFisher Scientific), following manufacturer’s instructions.

Standard chemical transformation was performed with MultiShotTM FlexPLate Mach1TM T1R (Invitrogen, ThermoFisher Scientific), grown O.N. at 37°C. Vector selection was performed with 100 μg/ml Spectinomycin (Sigma-Aldrich) in the growth medium for the pCR®8/GW/TOPO vectors, or 100 μg/ml Ampicillin (Normon) for the Z48 and ZED vectors. Plasmids were extracted with NZYMiniprep kit (NZYTech) and confirmed by Sanger sequencing, using M13 forward and M13 reverse primers for pCR®8/GW/TOPO vector, and GW reverse primer for Z48 and ZED vector. Final plasmids were purified with phenol/chloroform and concentration was determined by NanoDrop 1000 Spectrophotometer (ThermoFisher Scientific).

#### *In vitro* mRNA synthesis, Microinjection and Transgenesis

Z48 and ZED zebrafish lines were generated through TOL2-mediated transgenesis^38^. TOL2 cDNA was transcribed by Sp6 RNA polymerase (ThermoFisher Scientific) after Tol2-pCS2FA vector linearization with NotI restriction enzyme (Anza, Invitrogen, ThermoFisher Scientific). TOL2 mRNA was purified as previously described^36^. One-cell stage embryos were injected with 1nL solution containing 25 ng/µL of transposase mRNA, 25 ng/µL of phenol/chloroform purified plasmid (Z48 or ZED), and 0.05% phenol red.

#### Cas9 target design, sgRNA synthesis and mutant generation

Small guide RNAS (sgRNAs) targeting regions flanking zPtf1aE3 were designed using the Crisprscan software^39^ (SupplementaryTable16). Oligonucleotides (1,5μL at 100 μM each, from Sigma-Aldrich) were annealed *in vitro* by incubation at 95°C for 5 min in 2x Annealling Buffer (10mM Tris, pH7.5-8.0, 50mM NaCL, 1mM EDTA) followed by slow cooling at RT, and inserted into 100ng of pDR274 vector (#42250, Addgene) previously cut with BsaI (Anza, Invitrogen, ThermoFisher Scientific; 1:10). The pDR274 vectors carrying sgRNA sequences were linearized with HindIII, purified with phenol/chloroform and transcribed with T7 RNA polymerase (ThermoFisher Scientific). Final sgRNAs were purified as described previously^36^. One cell-stage zebrafish embryos were co-injected with two sgRNAs (40 ng/µl each) and Cas9 protein (300 ng/µl). Zebrafish mutant lines for zPtf1aE3 deletion were generated using the combinations sgRNA1+sgRNA2 and sgRNA3+sgRNA2 (SupplementaryTable16). Enhancer deletions in zebrafish were detected with PCR using HOT FIREPol DNA Polymerase (Solis BioDyne) with the flanking primers used to amplify the enhancers (SupplementaryTable16). PCR products were visualized by electrophoresis in 2% agarose gel and confirmed by Sanger sequencing. The mutations were further verified in the F1 mutants by sequencing.

#### Crispr-Cas9 in human cell lines

Four single-guide sequences named sg1, sg2, sg3, sg4, targeting hArid1abE3 enhancer were designed (SupplementaryTable16). sg1 and sg3 were designed upstream of the enhancer, while sg2 and sg4 were designed downstream of the enhancer. Two complementary oligonucleotides containing the single-guide sequences and BbsI ligation adapters were synthesized by Sigma. Two single-guide sequences for deletion of non-active region (based on H3K27ac), named ng1 and ng2, were used as negative control of the experiment ^20^. Oligonucleotides were annealed in T4 Ligation Buffer (ThermoFisher Scientific). sgRNA was cloned into the BbsI-linearized pSpCas9-T2A-GFP (#48138, Addgene) (sg1, sg3, ng1) and pU6-(BbsI)_CBh-Cas9-T2A-mCherry (#64324, Addgene) (sg2, sg4, ng2) vectors using T4 Ligase (ThermoFisher Scientific). The plasmid DNA was produced with Plasmid Midi Kit (Qiagen).

hTERT-HPNE cells were seeded in 6-well plates (1.1×10^5^ cells/well, at early passage number) and transfected (∼70-90% of confluency) using combinations: ng1+ng2; sg1+sg2; sg3+sg4. The transfection (1.5 µg of each sgRNA plasmid) was performed using lipofectamine 3000 (ThermoFisher Scientific), according to the manufacture instructions. Then, we changed to fresh culture medium after 24 h. Three independent replicates of the transfection were performed. After 48h of recovery, the cells were used in sub-sequent experiments.

#### Nucleic acid extraction from zebrafish and human cell lines

Genomic DNA was extracted from whole zebrafish embryos at 24 hpf, after removal of chorion, with the standard phenol/-chloroform DNA extraction, and used as template for PCR amplification in order to genotype the tested conditions (SupplementaryTable16). The DNA samples were resuspended in 20 μl of TE buffer with RNase [10mM Tris, pH 8.0; 1mM EDTA pH 8.0 and 100 μg/ml RNAse (Sigma-Aldrich)], incubated for 1 hour at 37°C, and stored at −20°C.

Genomic DNA from hTERT-HPNE cells was extracted 48h after transfection and used as template for PCR amplification in order to genotype the tested conditions (SupplementaryTable16).

RNA was extracted from zebrafish embryos with 500μl TRIzol (Invitrogen, ThermoScientific), following the manufacturer’s instructions. Samples were incubated 30min at 37°C with 1 μl DNAse I (ThermoScientific), 1μl 10x reaction buffer and 0.5μl NZY Ribonuclease Inhibitor (40U/μl) at 0.05μl/μl final concentration. After adding 1μl EDTA 50mM per 1μg of estimated RNA, final volume was completed to 60μl with H2O, phenol-chloroform standard purification was performed and the RNA stored at −80°C.

#### Immunohistochemistry in zebrafish embryos and human cell lines

Zebrafish embryos/larvae were euthanized by prolonged immersion in 200-300 mg/L tricaine (MS222; ethyl-3-aminobenzoate methanesulfonate, #E10521-10G, Sigma-Aldrich). Whenever necessary the chorion was removed, and the zebrafish were fixed in formaldehyde 4% (#F1635-500ML, Sigma-Aldrich) for 1h at RT (8-12dpf larvae) or O.N. at 4°C (48hpf embryos). Permeabilization was carried out by incubation with 1% Triton X-100 in PBS for 1h at RT, followed by blocking with 5% bovine serum albumin (BSA) in 0.1% Triton X-100 for 1h at RT. Zebrafish were incubated with the primary antibody diluted in blocking solution at 4°C O.N., and then incubated with the secondary antibody plus DAPI (1:1000, Invitrogen, ThermoFisher Scientific) diluted in blocking solution for 4 hours at RT. After each antibody incubation, embryos were washed 6 times in PBS-T (0.5 % Triton X-100 in PBS-1x) 5 minutes at RT. Embryos were stored in 50% Glycerol/PBS at 4°C before microscopy slides preparation in mounting medium (50% Glycerol/PBS). Images were acquired with confocal microscope Leica-SP5II (Leica Microsystems, Germany) and processed by ImageJ software. Primary antibodies: rabbit anti-Amylase (1:50, #A8273-1VL, Sigma-Aldrich), mouse anti-Alcam (1:50, #ZN-8, DSHB) and mouse anti-Nkx6.1 (1:50, #F55A10, DSHB). Secondary antibodies: goat anti-mouse AlexaFluor647 (1:800, #A-21236 Invitrogen, ThermoFisher Scientific), goat anti-rabbit AlexaFluor568 (1:800, #A-11036 Invitrogen, ThermoFisher Scientific).

The hTERT-HPNE cells were fixed at 48h after transfection in formaldehyde 4% (#F1635-500ML, Sigma-Aldrich) in PBS for 15 min at RT, permeabilized with 1% Triton X-100 in PBS and blocked with 2% BSA in PBS for 20 min at RT. Incubation with primary antibody in 2% BSA/PBS was O.N. at 4°C and in secondary antibody plus DAPI (1:1000, Invitrogen, ThermoFisher Scientific) was 3h at 4°C. in 2% BSA/PBS for 3h. Human cells were washed once after fixation and permeabilization, and three times after each incubation with primary and secondary antibodies with PBS 10 minutes at RT. Fluorescence images were obtained at 40x magnification on a Leica DMI6000 FFW. Primary antibody used: anti-ARID1A (1:1000; #HPA005456 Sigma-Aldrich). Secondary antibody used: donkey anti-rabbit (1:1000, #A31573, ThermoFisher Scientific).

#### Statistical Analysis

Two-tailed paired Student’s t-test from GraphPad Prism 5 (San Diego, CA, USA) was used. In all analyses, P<0.05 was required for statistical significance.

In hTERT-HPNE immunohistochemistry images, ARID1A nuclear staining was measured for each cell and normalized for the average staining of the nucleus of all the cells in the same field (ratio=ARID1A expression/mean of ARID1A expression in the field).

### II) Processing and Bioinformatic analysis

#### ChIP-seq analysis

Raw reads for the two replicates of high quality (FASTQC v.0.11.5^40^, Supplementary data 1 and 2) were aligned to the zebrafish genome (GRCz10/danRer10) using Bowtie2 (v.2.2.6) with default settings^41^. The file was converted into a bed file and the data extended 300 bp, bigwig tracks generated and uploaded to UCSC Genome Browser (Fig.1b). Highly enriched regions (peaks) were obtained by MACS14 (v.1.4.2) with the parameters “--nomodel, --nolambda and --space=30”^42^. Reproducibility of the two biological replicates was measured by Pearson’s correlation coefficient^43^.

#### Analysis of enhancer activity

To identify the best putative active enhancers, present in zebrafish adult pancreas, we intersected the peaks from the two H3K27ac replicates, selecting only the enriched regions present in both. Since H3K27ac is also present in promoter regions, we excluded peaks overlapping TSS coordinates. The genomic annotation of our putative enhancers was performed by HOMER annotatePeaks.pl^44^ (Fig. 1c). The adult pancreas putative active enhancer dataset (PsE+DevE) was crossed with the H3K27ac zebrafish embryonic dataset (dome, 80% epiboly, 24 hpf and 48 hpf) (SupplementaryTable15) ^33^ to identify enriched regions present only in adult pancreas (PsE) (Fig.1d). All genomic intersections were performed using Bedtools “intersect”^45^. We superimposed the H3K27ac mapped reads from adult pancreas and the embryonic dataset with the adult pancreas H3K27ac peaks using seqMINER (v1.3.4) with default settings (Fig.1d), showing read densities ±5 kb from the acetylation peak center^46^. Gene enrichment and functional annotation of our dataset were obtained with GREAT^15^, using the basal plus extension association rule (SupplementaryFig.3a).

#### ATAC-seq analysis

All libraries were sequenced on Illumina HiSeq 2500 platform and raw reads were mapped to the reference zebrafish genome (GRCz10/danRer10) using Bowtie2 (v.2.2.6) with parameters “-X 2000 and --very-sensitive”^41^. To avoid clonal artefacts, the duplicated mapped reads were removed using Samtools^47^. Mapped reads were filtered by the fragment size (≤120 bp) and mapping quality (≥10). For a better visualization, data were extended 10 bp, generated bigwig tracks and uploaded to the UCSC browser (Fig.1b). To call for enriched regions, MACS2 (v.2.1.0) ^42^ with the parameters “--nomodel, --keep-dup 1, --llocal 10000, --extsize 74, --shift – 37 and -p 0.07” were used. Reproducibility of the biological replicates was measured using the Pearson’s correlation coefficient^43^. Then, we applied the Irreproducible Discovery Rate (IDR) in order to obtain a confident and reproducible set of peaks^48^.

#### 4C-seq analysis

4C-seq libraries were first inspected for quality control using FASTQC (v.0.11.5^40^, Supplementary data 3-5) and demultiplexed using the script “demultiplex.py” from the FourCSeq package^49^, allowing for 1 mismatch in the primer sequence. A custom perl script was used for subsequent processing, as previously described^50,51^. Reads were aligned to the zebrafish genome (GRCz10/danRer10) using Bowtie^52^, keeping only uniquely mapping reads (v1.1.2, -m 1). Reads within fragments flanked by restriction sites of the same enzyme or smaller than 40 bp were filtered out. Mapped reads were then converted to reads-per-first-enzyme-fragment-end units, and smoothed using a 30 fragment mean running window algorithm (Fig. 4, 5).

#### HiChIP-seq analysis

HiChIP analysis from paired end fastq files to pairs of interacting fragments were performed using a custom python script based on the functions and defaults of the pytadbit python library^53^. This library first uses GEM mapper^54^ to map paired reads independently to the zebrafish reference genome (GRCz10/danRer10, flags used by GEM mapper --max-decoded-strata 1; --min-decoded-strata 0; -e 0.04). Then, reads are associated to a particular restriction fragment and paired together according to their read names. Once the reads are paired, the pairs of reads are filtered so that only those belonging to different restriction fragments are kept. Compressed sparse matrix files in cooler and hic formats were generated from those filtered reads using Cooler (“cload pairix” utility) and Juicer tools (“pre” utility) respectively for both visualization and further analysis. In order to predict the target promoter of putative enhancers a custom R script was designed, that uses uncompressed 5kb resolution sparse matrices as inputs. Those matrices were obtained using Juicer tools (“dump” utility) from the hic file and filtered for >=2 interactions between 2 interacting chunks < 100kb apart. In addition, only contacts connecting zebrafish pancreas active TSSs and putative active enhancers given by H3K27ac peaks from whole pancreas, adult pancreas (PsE), developing pancreas (DevE) and the different enhancer clusters (C1-C4) were included. This way, we could obtain the targets of putative active enhancers. An output table was produced with genes targeted by enhancers, per enhancer cluster (SupplementaryTable8). Custom scripts are provided in a gitlab repository (https://gitlab.com/rdacemel/pancreasregulome).

#### Identification of active promoters

H3K4me3 sequencing datasets (2 replicates performed in the HiChIP assay; Supplementary data 6-9) were aligned to the zebrafish genome (GRCz10/danRer10) using Bowtie2 (v.2.2.6) with default settings. Highly enriched regions (peaks) were obtained by MACS14 (v.1.4.2). algorithm with the parameters “--nomodel, -- nolambda and --space=30”^42^. Then, the peaks present in both replicates were filtered with the transcription start site (TSS) position to identify the active promoters using Bedtools “intersect”^45^.

#### RNA-seq analysis

Total RNA extracted from adult zebrafish (exocrine, endocrine and muscle) and sequenced on Illumina HiSeq 2000 platform was inspected for quality control using FASTQC (v.0.11.5^40^, Supplementary data 10-17). Then, sequences were trimmed to remove adaptors, sequencing artefacts and low-quality reads (Q<20)^55^. The BWA-MEM software was used to map the clean reads to the reference genome (ZV9/danRer7) with the parameters “-w 2 and -c 3”^56^. Gene expression was measured from the mapped reads using HT-seq-count ^57^. In addition, two public RNA-seq datasets were used (SupplementaryTable15).

#### Gene expression barplots

The average expression of genes associated with each enhancer cluster (PsE, DevE, C1-C4), as defined by HiChIP, was compared to the average expression of all genes present in the RNA-seq datasets using a custom R script and ggplot for drawing barplots (Fig.2a, SupplementaryFig.3b).

#### Conservation between zebrafish and human and PhastCons scores

To obtain the percentage of putative active enhancers conserved with human, the coordinates of putative active enhancers from adult zebrafish pancreas and embryos at different development stages (GRCz10/danRer10) were used as input to the UCSC genome coordinate conversion tool (https://genome.ucsc.edu/cgi-bin/hgLiftOver, liftover to hg19, October 2019) (Fig.3a). To visualize the conservation of the respective sequences, liftOver to hg38 was done and their average PhastCons conservation score plotted (Fig.3b). For this, we downloaded PhastCons scores in bigWig format from a 100-way multiple species alignment of vertebrates against human (hg38) (hg38.phastCons100way.bw, October 2019)^58^ and converted to BedGraph text format using the UCSC’s utility *bigWigToBedGraph*. Then, the Bedtools^45^ suite (v.2.27) was used to intersect and map different putative enhancer clusters in bed format with the conservation scores, storing for each putative enhancer the median and average PhastCons score. To know which of them overlap putative active enhancers in human pancreas, we used the Bedtools “intersect” tool with default ≥1 bp of overlap (Fig. 3b, blue). To check whether the conserved sequences showed a higher overlap with H3K27ac signal in humans than random, suggesting a higher likelihood of being also a putative active enhancer, we did the ratio of the number of conserved sequences overlapping H3K27ac signal in human over a similar one obtained from the average of a 10^5 random shuffling of the putative enhancer coordinates on the genome and their overlap with putative active enhancers in human (Fig. 3c).

#### Transcription factor binding motifs enrichment

To refine our data, H3K27ac peaks were filtered with the ATAC-seq peaks. Then, the transcription factor binding site (TFBS) predictor program Hypergeometric Optimization of Motif EnRichment (HOMER) was used to identify conserved sequence motifs enriched^44^. To evaluate our results, we also analysed, using HOMER, different acetylation data from: human pancreas, human ventricle, 24hpf and dome+80%epiboly (SupplementaryTable14 and 15). From each sample, the top 140 enriched motifs were selected, and the groups compared by performing hypergeometric enrichment tests (Fig. 3G, SupplementaryFig.7a-e). Chi-square test from GraphPad Prism 7 (v.7.04) was performed to evaluate the enrichment in 25 known pancreas-related TFs. The HOMER software was also applied in PsE, C1, C2, C3 and C4 in order to identify the TFBS present (SupplementaryFig.7f, SupplementaryTable14).

#### Identification of super-enhancers

We applied ROSE (Ranking Ordering of Super-Enhancers) algorithm with default parameters to define super-enhancers in our whole pancreas acetylation data and in human pancreas acetylation data^59^. Then, we performed gene ontology analysis in both data using PANTHER software (v.14.0, on April 2019) and compare the molecular functions obtained (http://pantherdb.org). To identify the genes shared between the two groups, we identified the human orthologous genes in our zebrafish list using Biomart (https://www.ensembl.org/biomart; on April 2019) and compared the groups (SupplementaryFig.6).

#### Disease association enrichment of genes from different enhancer clusters

Gene-disease association data were retrieved from DisGeNET (v.6.0) ^60^ (http://www.disgenet.org/), Integrative Biomedical Informatics Group GRIB/IMIM/UPF, on April 2019, selecting for pancreas-related diseases and filtering for a minimum score of 0.1 to exclude associations only from text-mining and keeping only diseases associated to at least 10 genes. Gene annotations were obtained from Ensembl via BioMart on April 2019 (BioMart export file available upon request) selecting protein coding genes in zebrafish and gene homologs between human and zebrafish. For each pancreas-related disease, zebrafish homologs of the human disease-associated genes were obtained and for the diseases remaining with at least 15 genes, hypergeometric tests were performed for each cluster (PsE, DevE, C1-C4) taking the set of zebrafish protein coding genes as the population size, the number of disease genes as number of successes and the number of genes in a given cluster as the sample size. The R package “qvalue” was used to correct for multiple testing using FDR and convert unadjusted p-values into q-values^61^. Hypergeometric enrichment was obtained as the ratio “(number disease genes in clusterX / number of genes in clusterX) / (number disease genes / number of protein coding genes)”. Finally, diseases with an absolute enrichment ≥ 1.5 and a q-value ≤ 0.05 were considered significantly enriched(/depleted) in the respective cluster (Fig.3d).

## Table of contents

### List of supplementary tables

**Supplementary table 1** – Genomic coordinates of pancreatic enhancers (excel file)

**Supplementary table 2** – List of genes (excel file)

**Supplementary table 3** – List of active promoters captured by HiChIP (H3K4me3) (excel file)

**Supplementary table 4** – List of putative enhancers (PsE, DevE, Embryo) (excel file)

**Supplementary table 5** – List of putative enhancers (C1-C4) (excel file)

**Supplementary table 6** – Enrichment of the expression of genes associated by GREAT to PsE (Output from GREAT) (excel file)

**Supplementary table 7** – Enrichment of the expression of genes associated by GREAT to DevE (Output from GREAT) (excel file)

**Supplementary table 8** – HiChIP results: number of interactions between a 5kb bin containing a putative active enhancer in whole pancreas and a 5kb bin containing the TSS of the indicated gene (excel file)

**Supplementary table 9** – Zebrafish tissue RNA-seq raw counts (excel file)

**Supplementary table 10** – List of putative pancreatic enhancer sequences tested (excel file)

**Supplementary table 11** – Fig3C stats (100k times shuffling hg38 coordinates lifted over from zebrafish putative active enhancer sets and checking their intersect with human pancreas H3K27ac peaks) (excel file)

**Supplementary table 12** – Hypergeometric test enrichment for the association to pancreas-related diseases on the genes from each enhancer cluster (Embryo, C1-C4, PsE) (excel file)

**Supplementary table 13** – Super-enhancers active in zebrafish pancreas predicted by ROSE (excel file)

**Supplementary table 14** – List of TF found by HOMER (excel file)

**Supplementary table 15** – List of data used (excel file)

**Supplementary table 16** – List of primers used (excel file)

### List of supplementary data

Supplementary data 1 – Pancreas H3K27ac ChIP-seq fastqc 1 (html file)

Supplementary data 2 – Pancreas H3K27ac ChIP-seq fastqc 2 (html file)

Supplementary data 3 – 4C-seq Arid1ab fastqc 1 (html file)

Supplementary data 4 – 4C-seq Arid1ab fastqc 2 (html file)

Supplementary data 5 – 4C-seq Ptf1a fastqc (html file)

Supplementary data 6– Pancreas H3K4me3 HiChIP fastqc 1-1 (html file)

Supplementary data 7– Pancreas H3K4me3 HiChIP fastqc 1-2 (html file)

Supplementary data 8– Pancreas H3K4me3 HiChIP fastqc 2-1 (html file)

Supplementary data 9– Pancreas H3K4me3 HiChIP fastqc 2-2 (html file)

Supplementary data 10– RNA-seq Endocrine old fastqc 1 (html file)

Supplementary data 11– RNA-seq Endocrine old fastqc 2 (html file)

Supplementary data 12– RNA-seq Endocrine young fastqc 1 (html file)

Supplementary data 13– RNA-seq Endocrine young fastqc 2 (html file)

Supplementary data 14– RNA-seq Exocrine old fastqc (html file)

Supplementary data 15– RNA-seq Exocrine young fastqc (html file)

Supplementary data 16– RNA-seq Muscle young fastqc (html file)

Supplementary data 17– RNA-seq Muscle old fastqc (html file)

**Supplementary figure 1.**
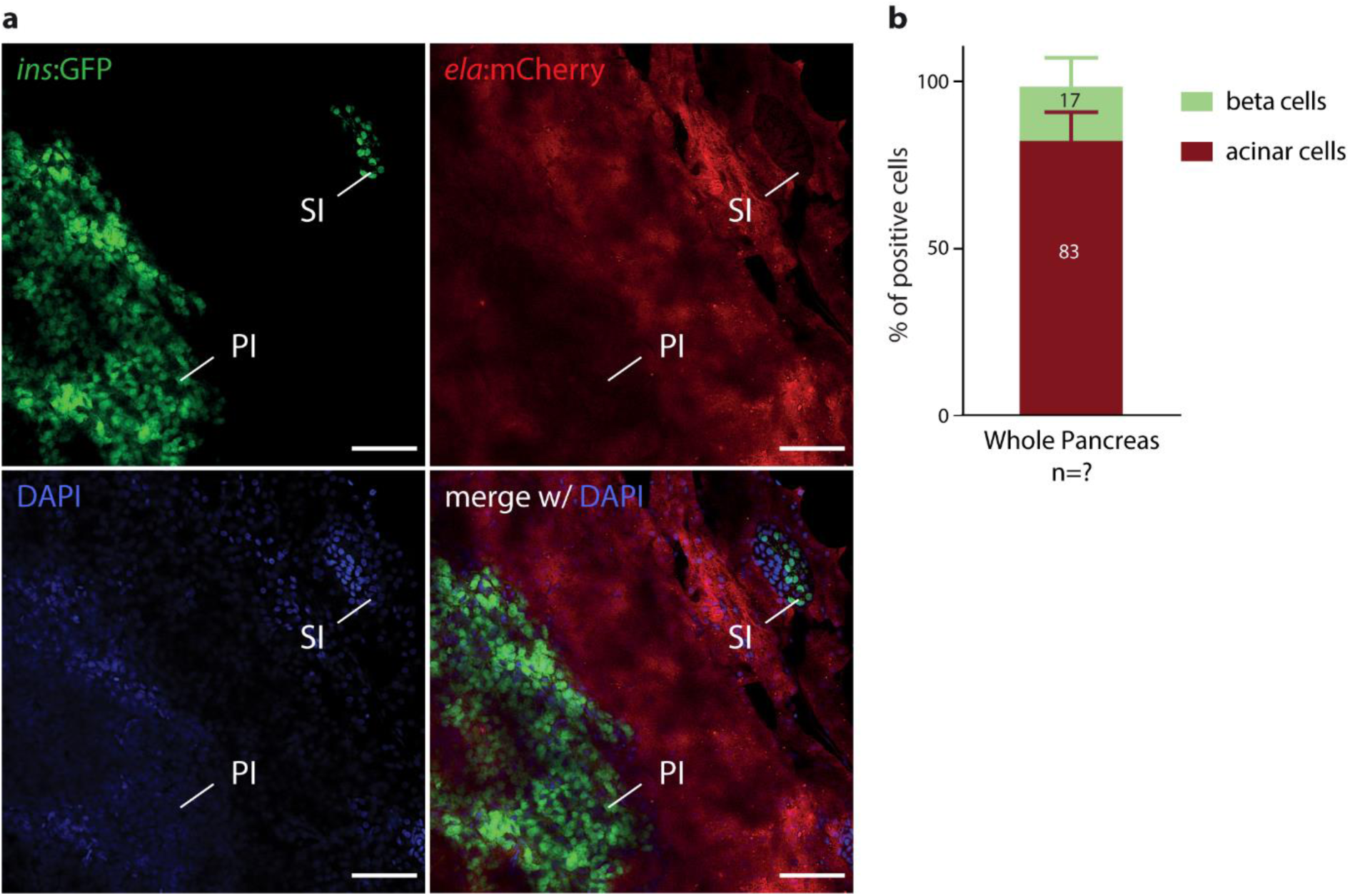
Basic constitution of the zebrafish adult pancreas. **a)** Confocal image of whole-mounted pancreatic tissue from adult Tg(insulin:GFP, elastase:mCherry) zebrafish. The endocrine pancreas consists of a larger principal islet (PI) and smaller secondary islets (SI), where the insulin-producing beta-cells are visible (green), embedded within the pancreatic exocrine tissue, that itself is comprised of acinar cells (red) and a network of duct cells. Nuclei are stained with DAPI. Scale bar: 50 μm. **b)** Percentage of pancreatic endocrine beta-cells and exocrine acinar cells from Tg(insulin:GFP, elastase:mCherry), quantified by flow cytometry, from adult zebrafish pancreas (n=5).

**Supplementary figure 2.**
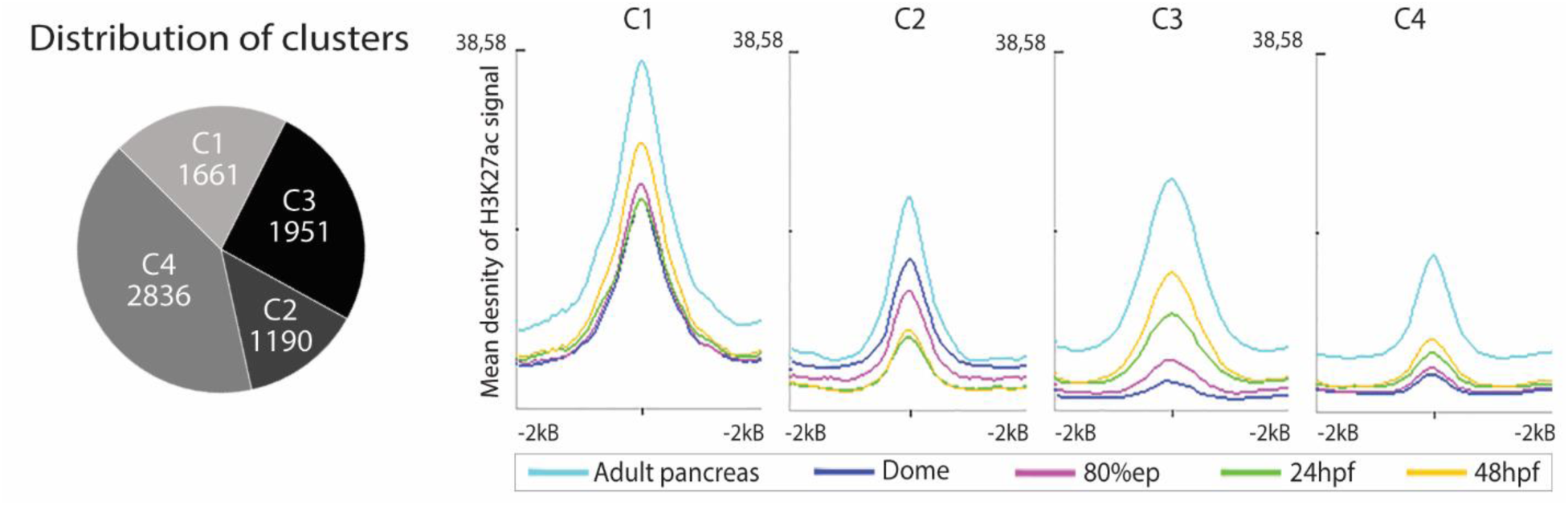
Left panel represents the number of sequences contained in each of the four clusters observed in DevE; C1: Cluster 1, C2: Cluster 2, C3: Cluster 3 and C4: Cluster 4. Right panel represents the mean density of H3K27Ac signal for each cluster, centred in its summit and expanded ±2kb, from adult pancreas and embryos at different developmental stages: dome, 80% epiboly, 24hpf and 48hpf.

**Supplementary figure 3.**
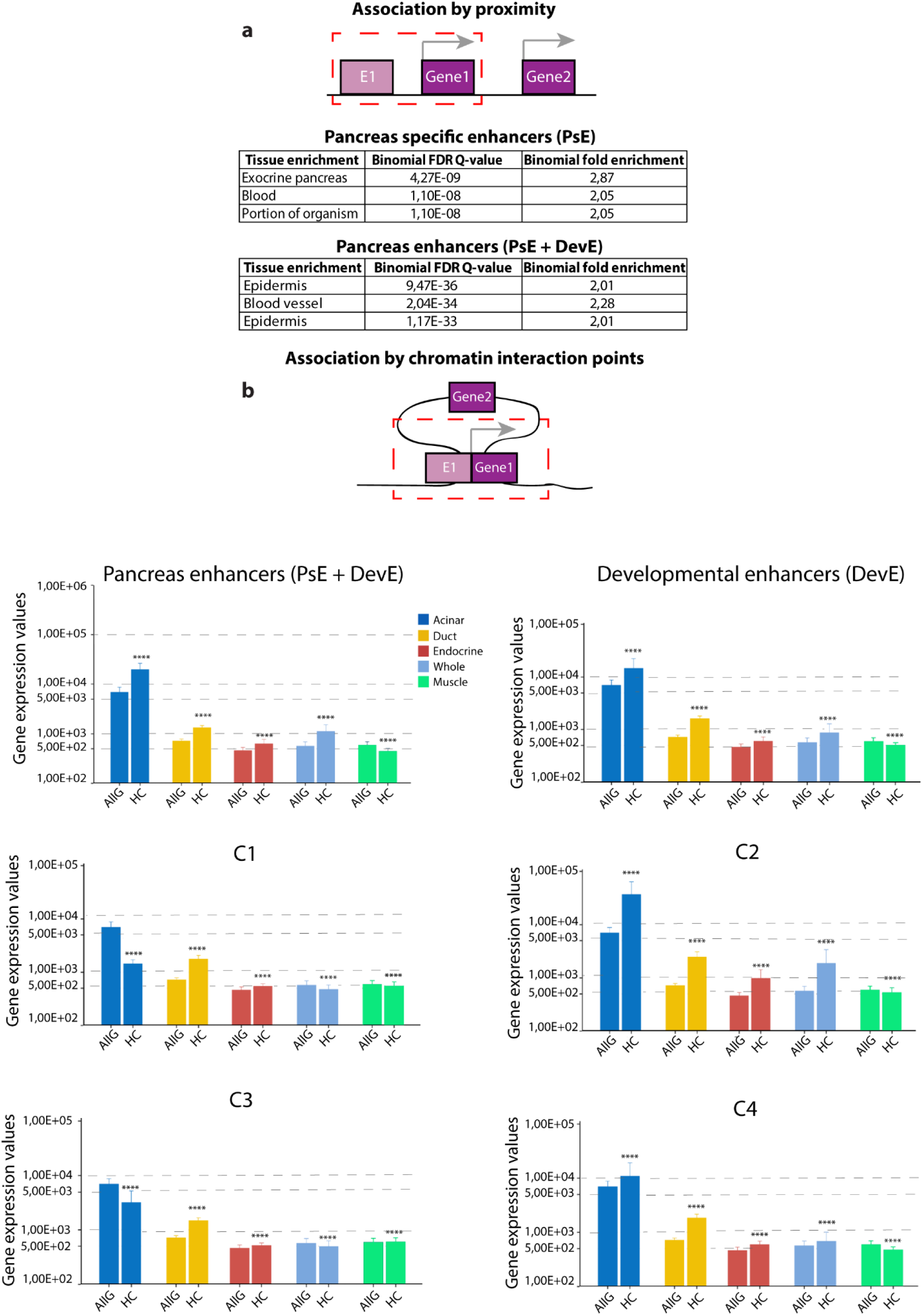
a) Upper panel: Schematic representation of gene-to-enhancer association by genomic proximity with GREAT^1^. Lower panel: Tissue specific expression enrichment for genes associated to PsE and PsE+DevE. **b)** Schematic representation of gene-to-enhancer association by chromatin interaction points defined by HiChIP for H3K4me3 (HC). Average expression of genes interacting with different enhancer sets detected by HC in adult zebrafish pancreas, compared to the average expression of all genes (AllG), determined by RNA-seq from pancreatic cells (acinar, duct, endocrine), whole pancreas and muscle (control). Enhancer sets: pancreas specific enhancers (PsE), developmental enhancers (DevE), and clusters of developmental enhancers C1, C2, C3 and C4. Wilcoxon test, *p*-values ≤ 0.05 considered significant. PsE+DevE:****p<2e-16; DevE, C1-C4:\*\*\*\**p*<0,0001.

**Supplementary figure 4.**
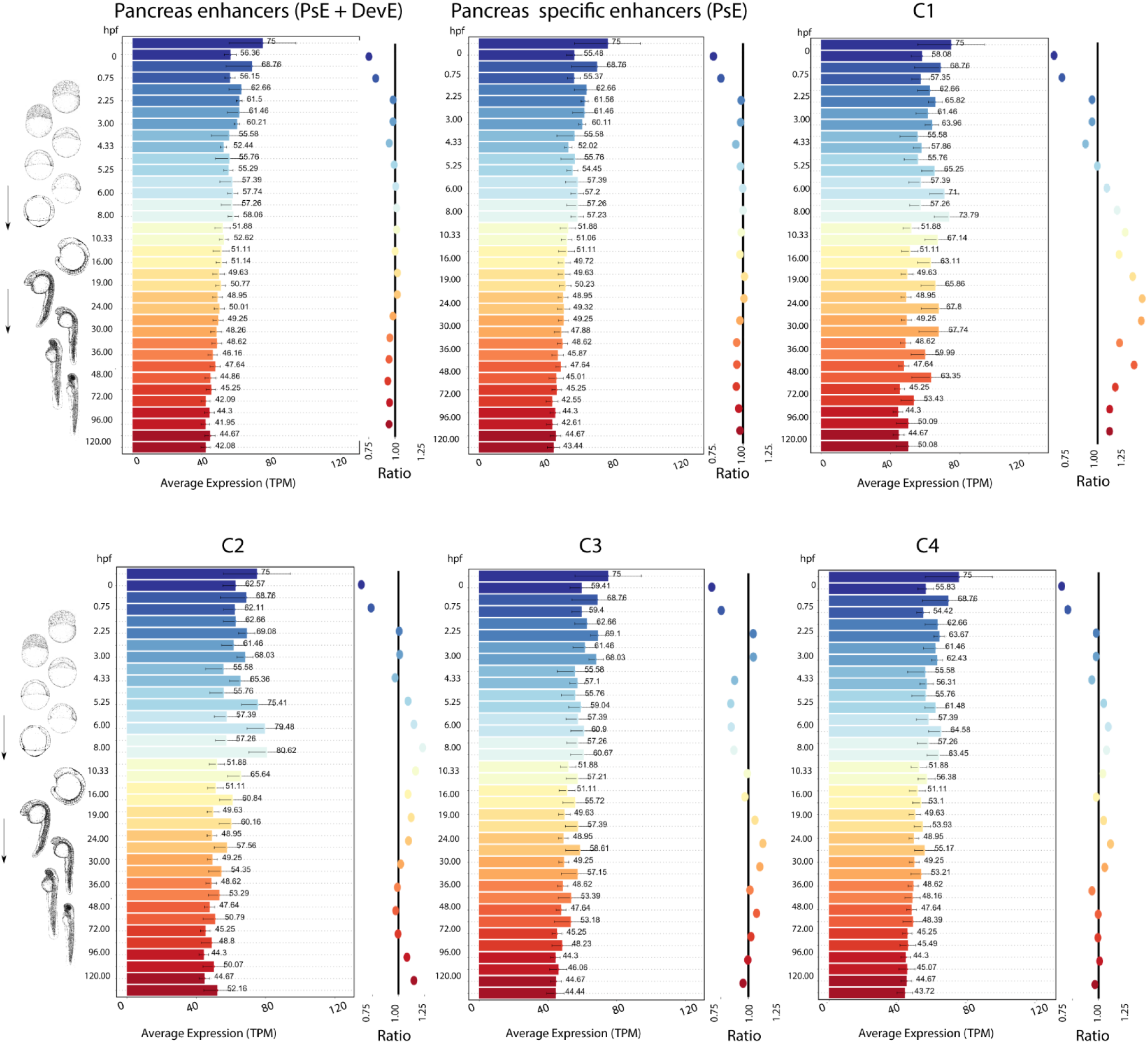
At the left side of each panel, the average gene expression (transcripts per million, TPM) detected from RNAseq from zebrafish embryos at different developmental stages (0 to 120hpf) ^2^ is plotted. The top bar of each colour is the average expression of genes associated to pancreas enhancers by HiChIP and bottom bar of each colour is the average expression of all genes as control, with the respective value depicted for each bar. On the right side of each panel is the depicted the ratio of the average expression of all genes as control by the average expression of genes associated to pancreas enhancers by HiChIP, maintaining the same colour code.

**Supplementary figure 5.**
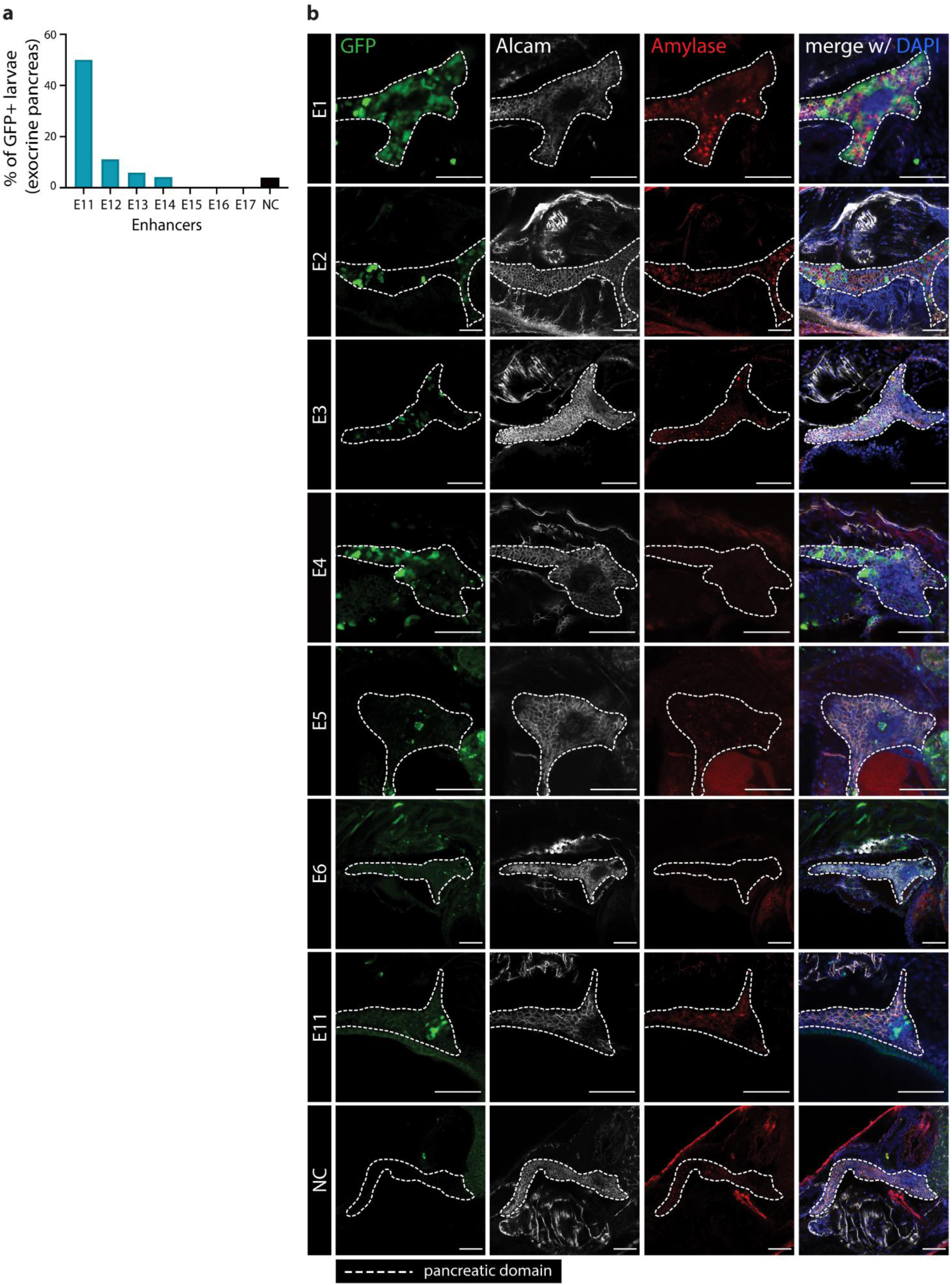
*In vivo* validation of selected putative pancreatic enhancers, identified by ATAC and ChIP-seq data, by transgenesis assays in zebrafish embryos injected with Z48 vector carrying putative enhancer sequences and the empty vector was used as negative control (NC). **a)** Percentage of embryos with GFP expression in zebrafish pancreas for sequences (E11 to 17) with low H3K27ac ChIP signal value: ((−log10(p-value)<35). Chi-square test, p-values<0.05 were considered significant. NC is the negative control (empty vector). **b)** Representative confocal images of transgenic F0 larvae for all validated enhancers (E1-E6 and E11); whole mount immunohistochemistry of 10-12 dpf zebrafish larvae showing enhancer activity, in the form of GFP expression (green), within the pancreatic domain (dashed white line), in comparison to the NC. The exocrine pancreas is stained with anti-Alcam (white) and anti-Amylase (red) antibodies. Nuclei are stained with DAPI. Scale bar, 60 μm.

**Supplementary figure 6.**
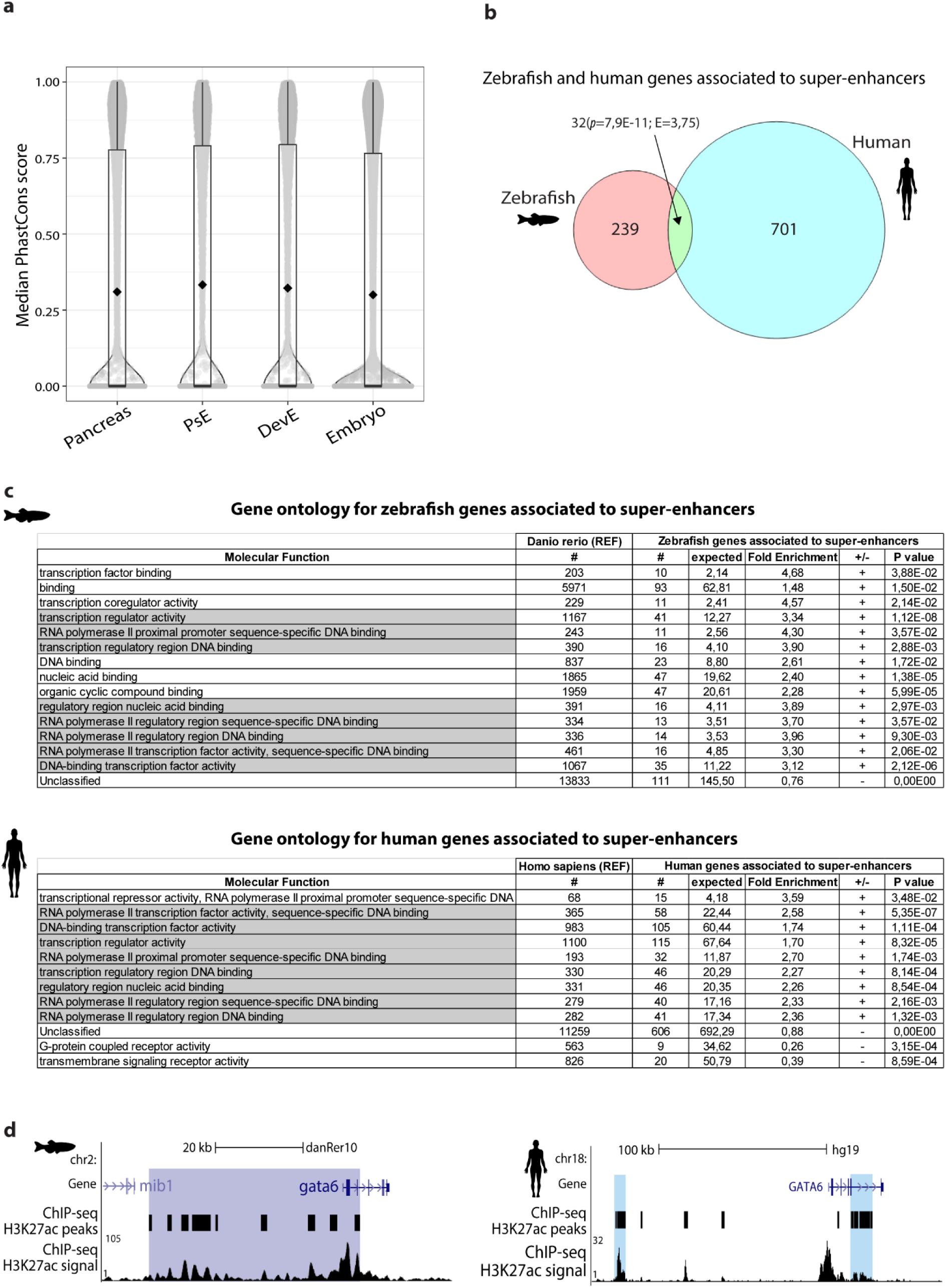
**a)** Distribution of the median PhastCons scores for each zebrafish putative enhancer sequence active in adult pancreas (14301), adult pancreas only (PsE, 6918), adult pancreas and embryo (DevE, 8368) and in embryo (Embryo, 65871). Putative active enhancer sequences converted from DanRer10 to DanRer7 (liftOver) to match the available conservation scores for zebrafish and 7 vertebrates. The median value is zero for the 4 enhancer sets (lower bar of the boxplots) since most sequences are not conserved, while the average is, respectively, 0.31, 0.33, 0.32, 0.30 (back diamond inside the boxplot). **b)** Zebrafish and human genes associated to super-enhancers, identified by ROSE^3^ **c)** Gene ontology for genes associated to super-enhancers in zebrafish (upper table) and human (lower table). **d)** H3k27ac profile of the landscape of a gene important in pancreatic development in zebrafish (gata6; left) and human (GATA6; right); super-enhancers are highlighted in purple (zebrafish) and blue (human).

**Supplementary figure 7.**
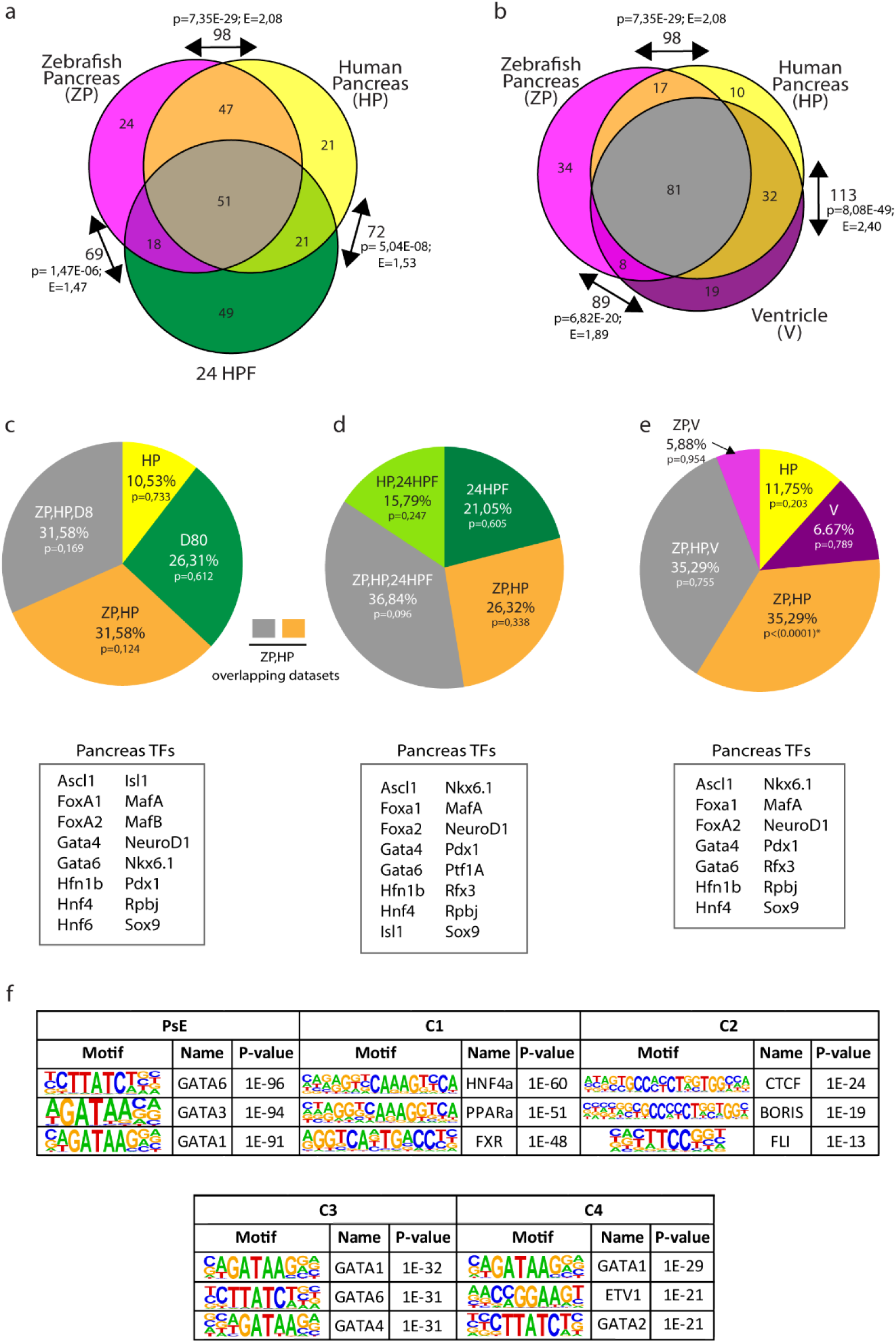
**a-b)** Regions enriched in ATAC-seq and H3k27ac signal from zebrafish pancreas (ZP), human pancreas (HP), 24hpf zebrafish embryos (24HPF) and human ventricle (V) were investigated for TFBS motifs (SupplementaryTable14 and 15). The top 140 enriched TFBS motifs from each tissue were selected and the overlap of those sets was analysed. Arrows: number of TFBS motifs shared between two different groups, the enrichment of TFBS motifs and respective *p*-value for each intersection. Number of TFBS motifs identified in each group/intersection. **a)** ZP, HP and 24HPF; **b)** ZP, HP, V. **c-e)** The motifs corresponding to 25 pancreas transcription factors (Pancreas TFs) selected from literature were analysed for their representation in H3K27ac peaks filtered with ATAC-seq peaks among the different zebrafish (ZP, 24HPF, D80) and human tissues (HP, V), and tissue-intersections shown in a-b). The distribution of these TFBS motifs through the different tissues is shown in the graph with the percentage and the respective *p*-value indicated for each group (top). The TFs present in the tissues, from each graph, are depicted in the table (bottom). **c)** ZP, HP and dome+80%epiboly (D80); **d)** ZP, HP and 24hpf; **e)** ZP, HP and V. Statistical significance was assessed by Chi-square test, *p*-values<0.05 were considered significant. **f)** List of top three TFBS motifs enriched in H3K27ac ChIP-seq data, for the different enhancer sets: pancreas specific enhancers (PsE) and clusters of developmental enhancers C1, C2, C3 and C4, with the respective *p*-value.

**Supplementary figure 8.**
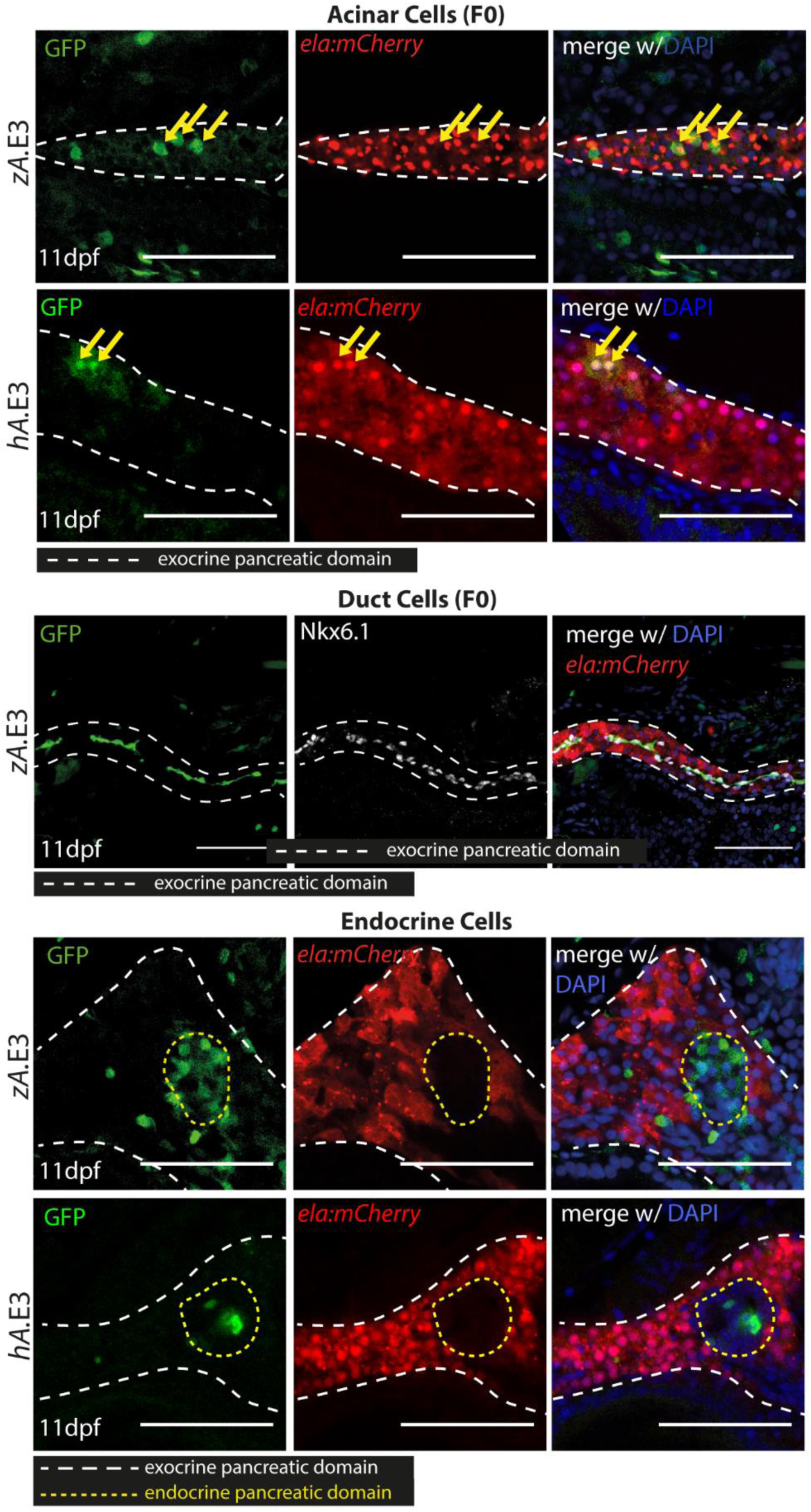
Human and zebrafish *ARID1A/arid1ab* enhancers drive expression in various pancreatic cell types. Confocal images of zebrafish pancreas (white dashed line), at 11dpf, from F0 Tg(*ela:mCherry*) embryos injected with Z48 vector carrying zebrafish zAridE3 (zA.E3) or human hAridE3 (hA.E3) enhancers. The top panel shows elastase-producing acinar cells (red), the middle panel shows NKX6.1-positive duct cells (white), and the bottom panel shows the endocrine pancreas domain (yellow dashed line). In all panels, the exocrine pancreas domain is indicated with a white dashed line and nuclei are stained with DAPI (blue). Scale bar: 60 µm.

**Supplementary figure 9.**
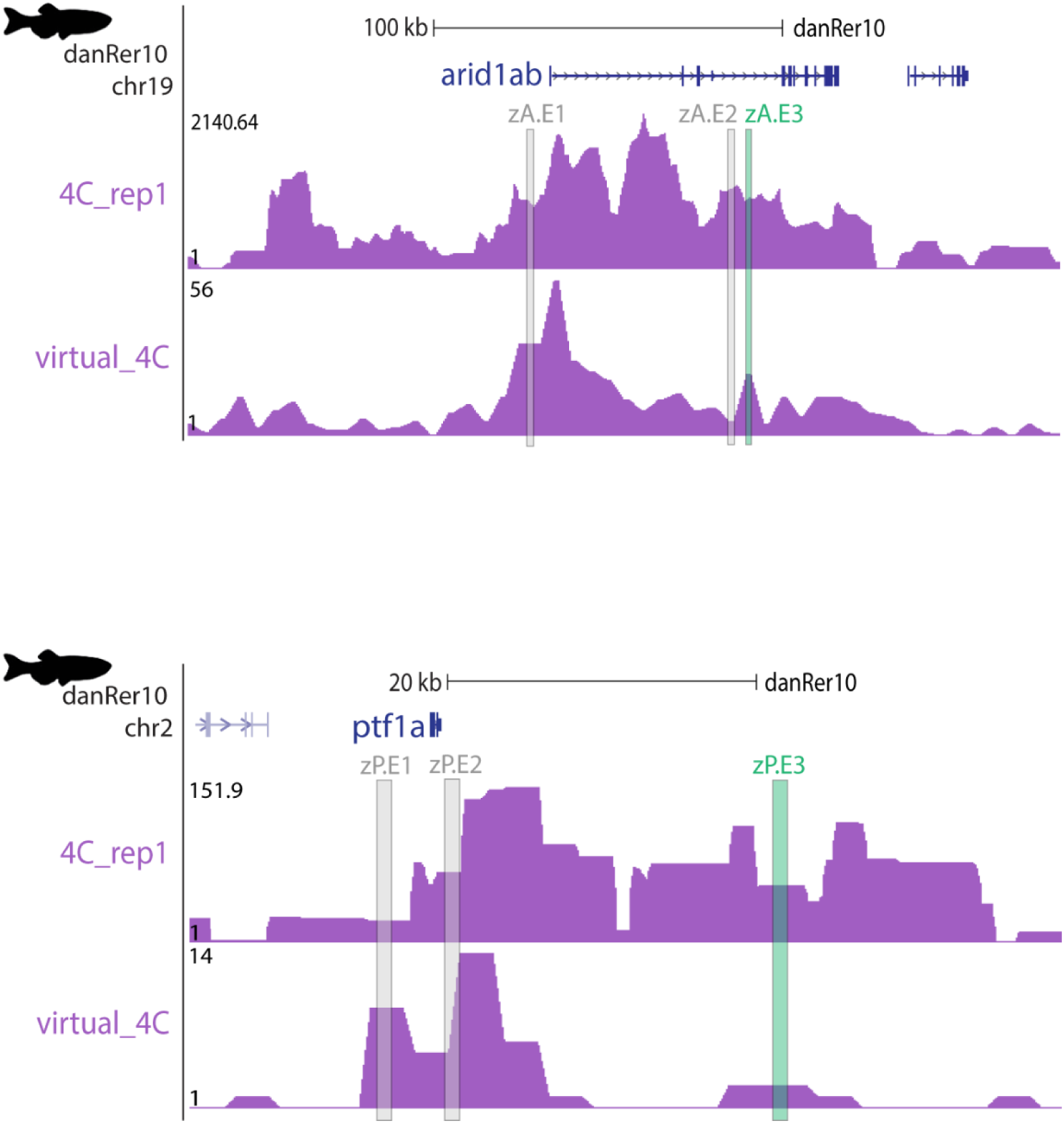
Genomic landscape of zebrafish *arid1ab* gene (top) and *ptf1a* gene (bottom), showing *arid1ab* 4C interactions (purple) from 4C-seq replicates and virtual 4C from HiChIP for H3K4me3 in adult zebrafish pancreas. Tested putative enhancers are highlighted in grey, zA.E1 (zAridE1) and zA.E2 (zAridE2) and green zA.E3 (zAridE3).

**Supplementary figure 10.**
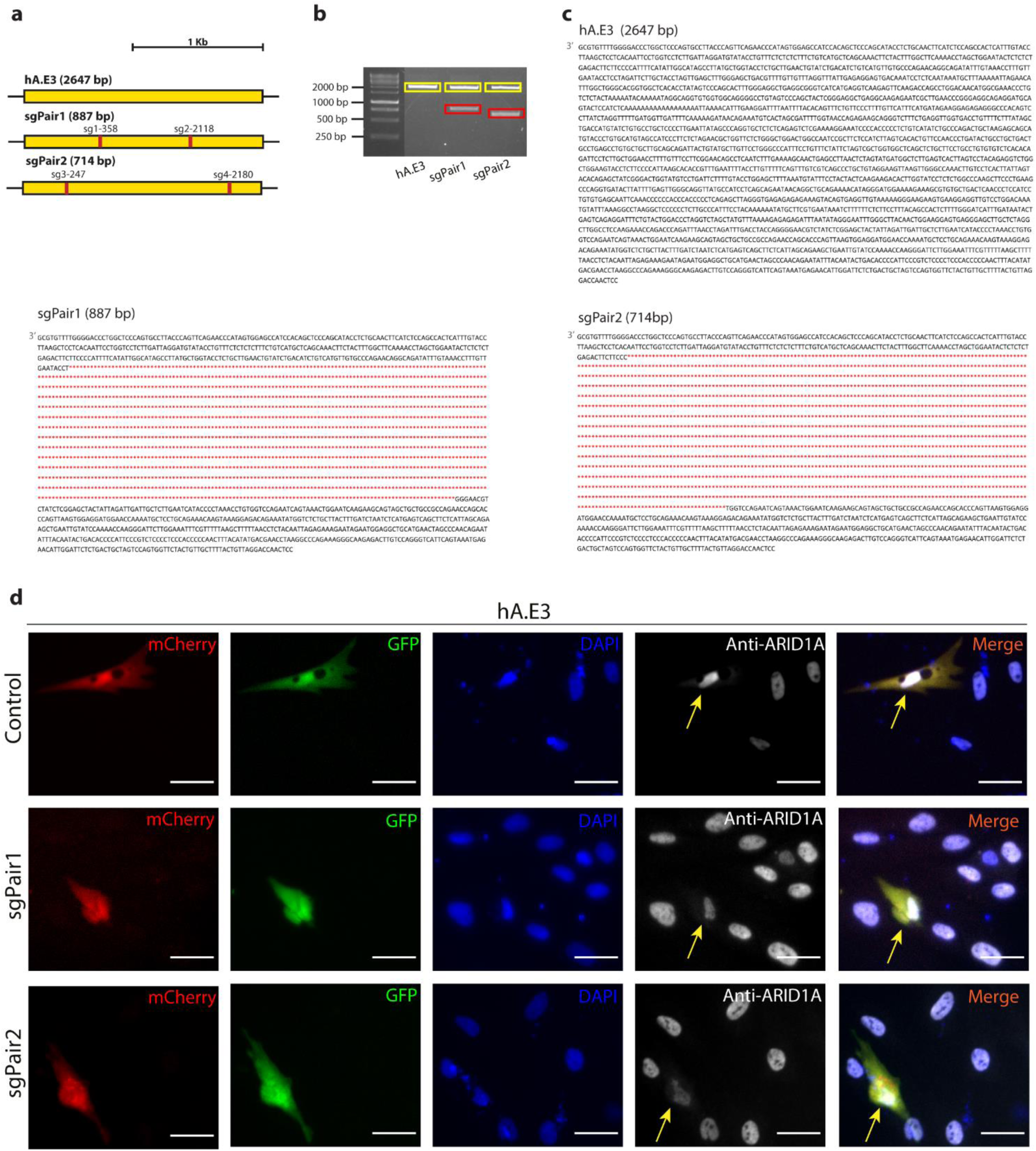
**a)** Schematic depiction of the targeting strategy for deletion of the hA.E3 (hAridE3) locus, indicating the CRISPR sgRNA target sites (red). **b)** Agarose gel showing the wild-type (yellow) and deleted (red) PCR amplified hA.E3 sequence after gene editing with the respective sgRNA pair. **c)** DNA sequencing results of the indicated genomic PCR products (yellow and red boxes). **d)** Representative fluorescent microscopy images of transfected hTERT-HPNE human cells, defined by the co-expression of GFP and mCherry (yellow arrows). Nuclei were stained with DAPI (blue) and anti-ARID1A antibody (white). The yellow arrows are pointing to the double transfected cell. Images were captured with Leica DMI6000 FFW microscope. Scale bar: 40 μm.

**Supplementary figure 11.**
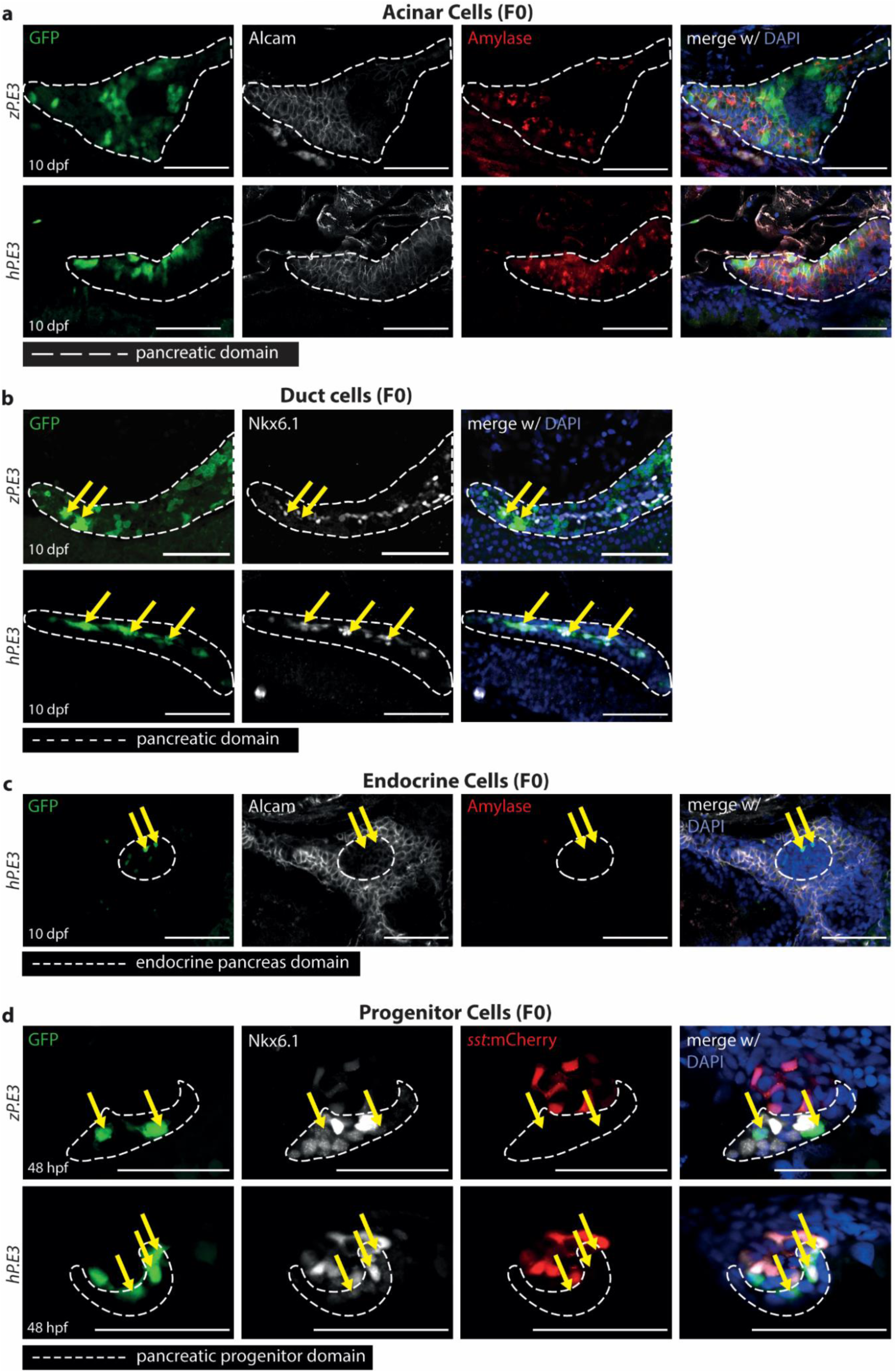
Human and zebrafish distal *PTF1a/ptf1a* enhancers drive similar reporter expression in various pancreatic cell types. **a-d)** Representative confocal images of GFP expression (green) driven by zebrafish zP.E3(zptf1aE3) or human hP.E3 (hptf1aE3) sequences in F0 10dpf zebrafish larvae in **a)** pancreatic acinar cells, stained with anti-Alcam (white) and anti-Amylase (red), **b)** duct cells, stained with anti-NKX6.1 (white) and **c)** endocrine cells, delimited by absence of staining of acinar cell markers. **d)** Pancreatic progenitor cells of 48hpf F0 zebrafish embryos stained with anti-NKX6.1 (white), adjacent to the endocrine principal islet identified by somatostatin expression in differentiated delta-cells, using Tg(somatostatin:mCherry) zebrafish. Nuclei are stained with DAPI (blue). Yellow arrows point to localization of GFP within the respective identified tissues (white dashed line). Scale bar 60 μm.

**Supplementary Figure 12.**
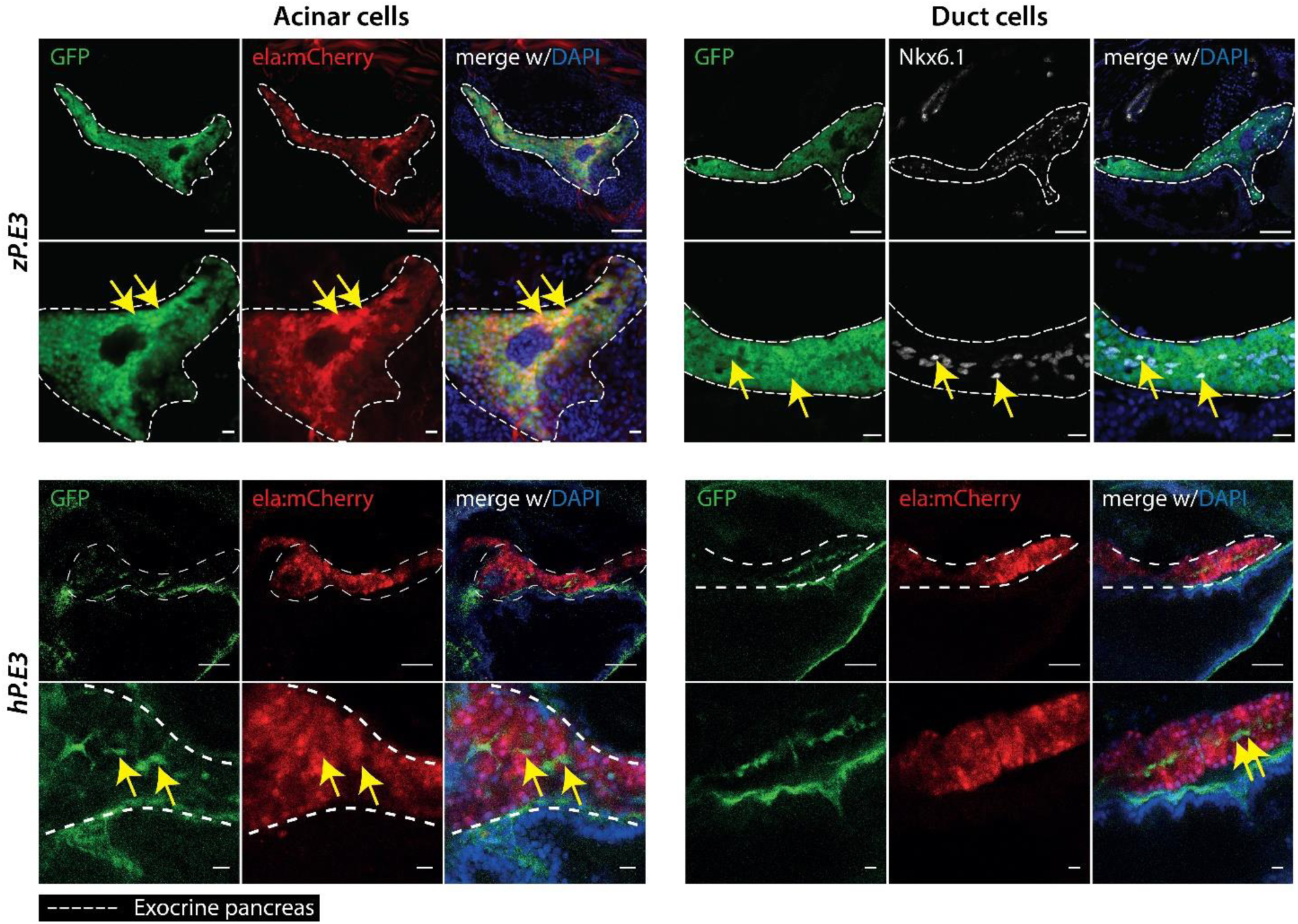
Zebrafish and human distal *ptf1a/PTF1a* reporter transgenic lines, zP.E3 (zptf1aE3) and hP.E3 (hptf1aE3), respectively, drive GFP expression in pancreatic acinar cells and duct cells. Left: Representative confocal images of a double zebrafish reporter transgenic line (zE3:GFP and elastase:mCherry) and a human reporter transgenic line (hP.E3: GFP), showing zP.E3 and hP.E3 mediated GFP expression (green) co-localizing with the exocrine specific elastase mCherry mediated expression (red). Right: Representative confocal images of a double zebrafish reporter transgenic line (zE3:GFP and elastase:mCherry) co-localizing with Nkx6.1, a specific duct cell marker, and for the human reporter transgenic line (hP.E3: GFP), showing hP.E3 mediated GFP expression (green) not co-localizing with elastase:mCherry. 8dpf larvae were labeled with Nkx6.1 (white) and nuclei were labeled with DAPI (blue). Yellow arrows point to examples of GFP and pancreatic markers co-localization. Scale bar 60 μm.

**Supplementary figure 13.**
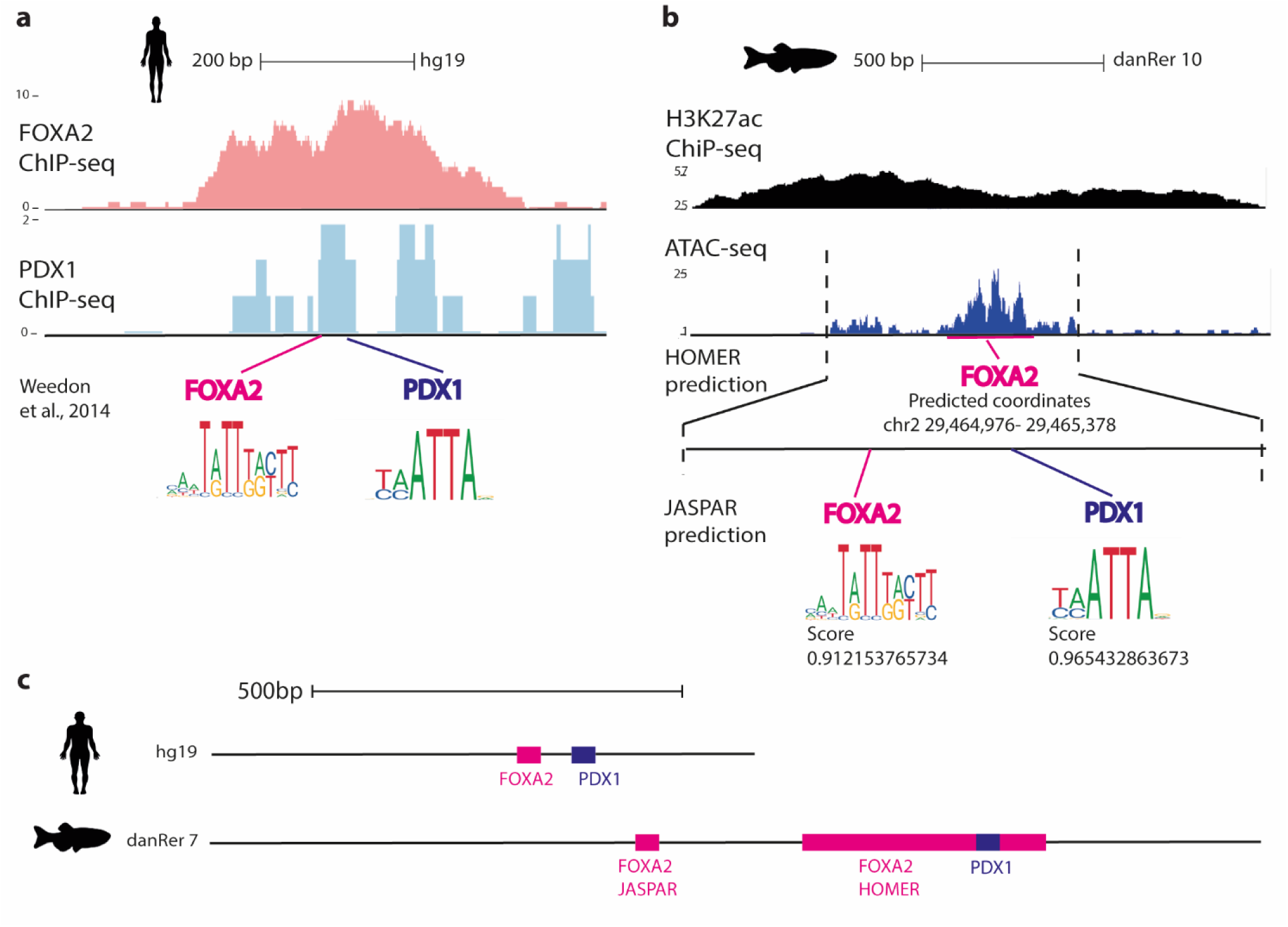
**a)** ChIP-seq density plots at the hP.E3 (hptf1aE3) locus showing FOXA2 (pink) and PDX1 (blue) ChIP-seq peaks generated from human endocrine islets^4^. The location of the respective predicted binding sites (verified by Weedon et al., 2014)^5^ are depicted below. b) H3K27ac ChIP-seq (black) and ATAC-seq (blue) read density plots at the zP.E3 (zptf1aE3) locus, and putative FOXA2 and PDX1 transcription factor binding sites predicted by JASPAR^6^ with respective score. c) Schematic representation of the human and zebrafish ortholog enhancers; hP.E3 (above) and zP.E3 (below).

**Supplementary figure 14.**
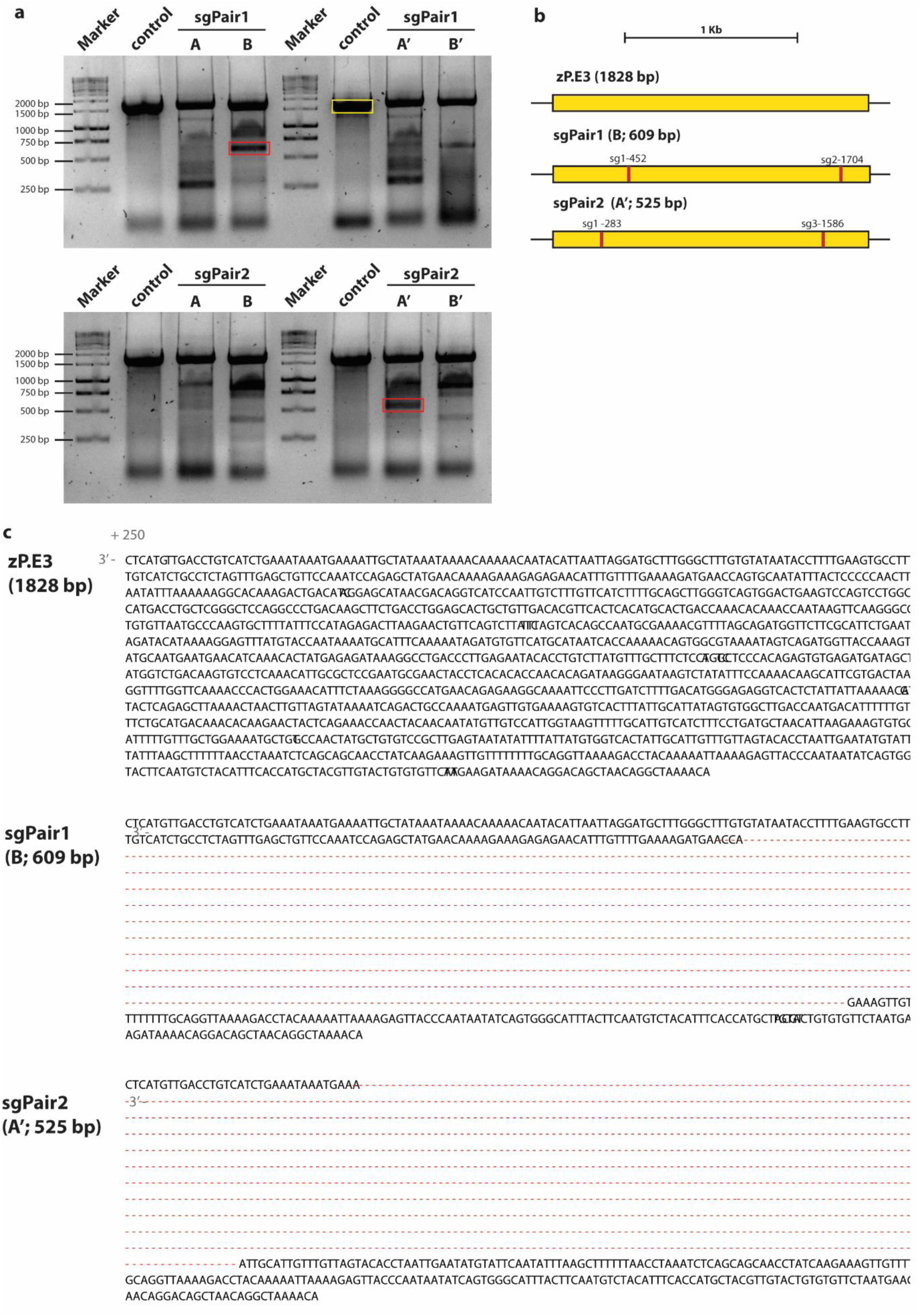
CRISPR/Cas9-mediated deletion of the zP.E3 (zptf1aE3) enhancer in F0 zebrafish. **a)** PCR screening of zptf1aE3 deletion after co-injection of zebrafish embryos with Cas9 and different combinations of sgRNAs: successful deletions appear as truncated PCR products (red rectangle), in comparison with the wild-type sequence from non-injected embryos (control, yellow rectangle). **b)** Schematic representation of the deletions induced by CRISPR/Cas9. **c)** Sequence of the wild-type and representative sequences of enhancer deletion, corresponding to the annotated red rectangles in the gel (a).

**Supplementary figure 15.**
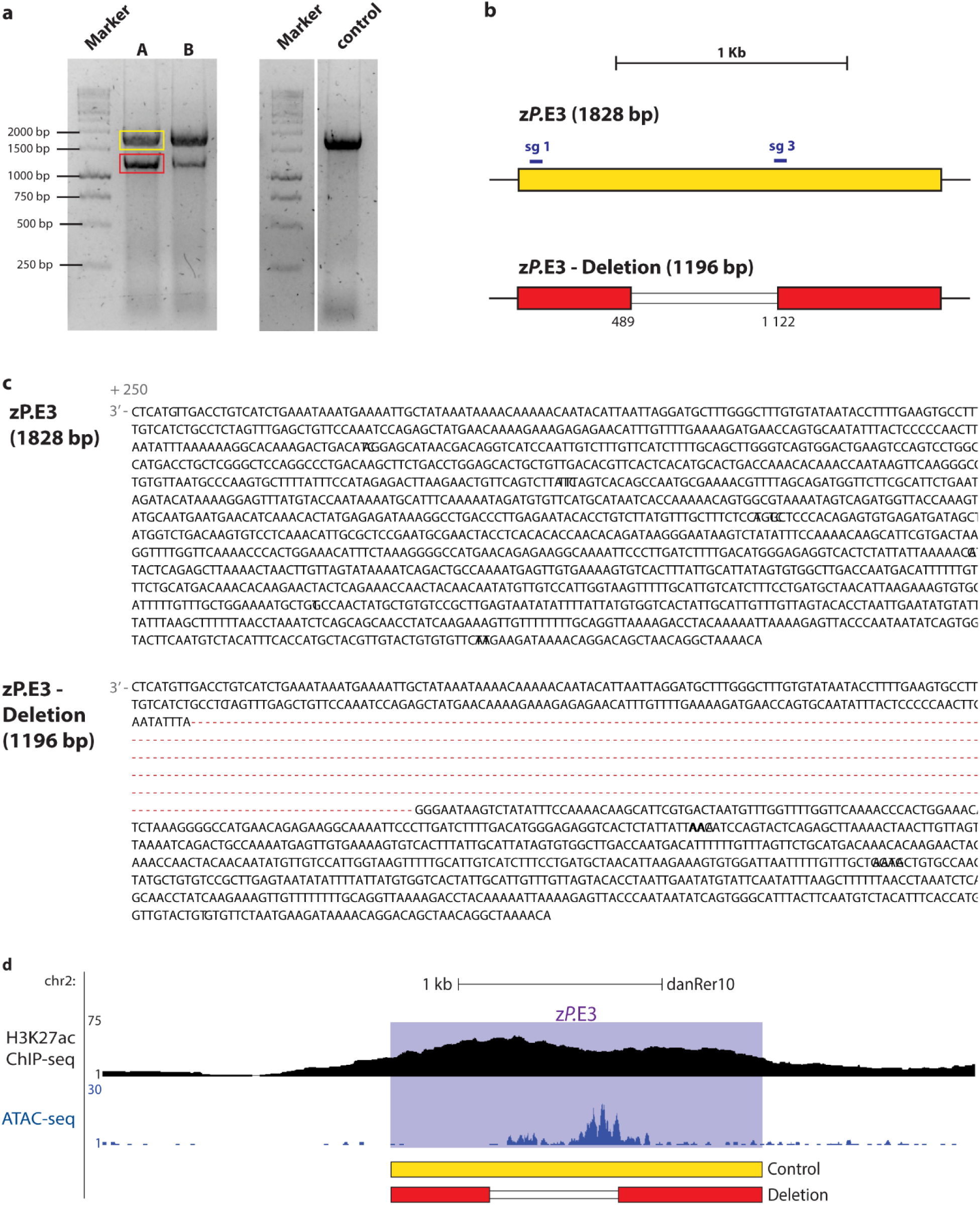
Mutant zebrafish with zP.E3 (zptf1aE3) enhancer deletion. **a)** PCR screening of genomic deletions for zP.E3 enhancer in zebrafish embryos, appearing as truncated PCR products (red rectangle), in comparison with the wild-type sequence from the same batch siblings (control; yellow rectangle). **b)** Schematic representation of the 632 bp deletion induced by CRISPR/Cas9. **c)** DNA sequencing results of the indicated genomic PCR products (yellow and red rectangles). **d)** Alignment of the wild-type and deleted alleles to the zebrafish, genome showing the ATAC-seq and ChIP-seq profiles at the zP.E3 locus.

**Supplementary figure16.**
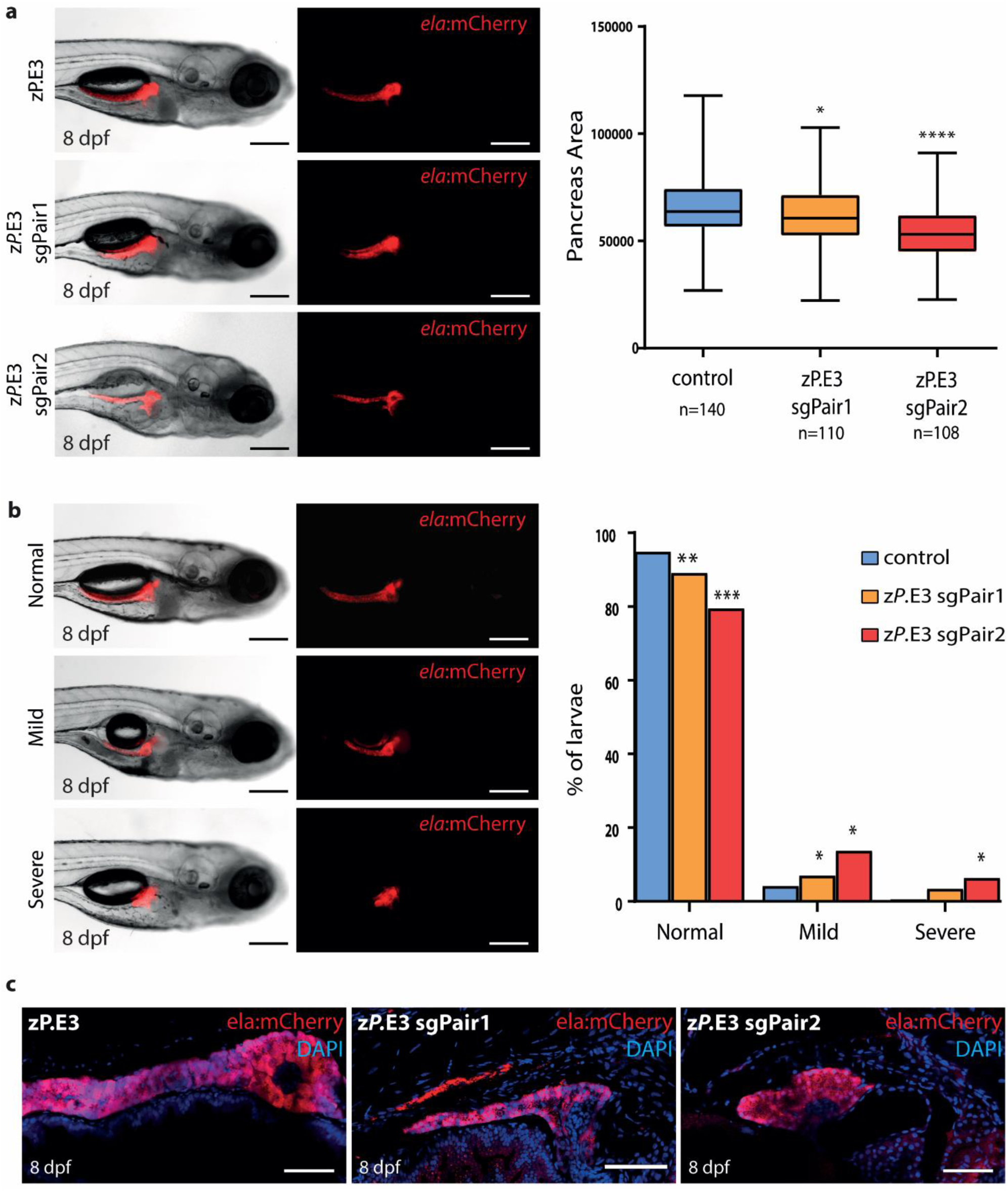
CRISPR/Cas9-mediated deletion of zP.E3 (zptf1aE3) affects pancreas size in F0 zebrafish. **a)** Tg(*elastase:mCherry*) embryos were injected with Cas9 alone (control) or co-injected with Cas9 and a pair of sgRNAs (zP.E3 sgPair1 or sgPair2) and monitored at 8 dpf. Representative live images are shown in the left panels. Scale bar: 250 µm. Quantification of total pancreas area are represented in the right panels. Unpaired student’s t-test (two-tailed), p-values<0.05 were considered significant (****p<0.0001, *p<0,05). **b)** Larvae where phenotyped as either normal or as one of the two following classes: mild pancreatic defect with significantly reduced pancreas size (mild) and severe pancreatic defects with reduced pancreas with abnormal structure (severe). Fisher’s exact test, p-values<0.05 were considered significant (***p<0.001, **p<0.01, *p<0.05). **c)** Representative confocal fluorescent images of whole-mounted 8 dpf Tg(*elastase:mCherry*) larvae showing impaired development of pancreas upon CRISPR/Cas9-mediated deletions of the zP.E3 enhancer (zP.E3 sgPair1 and zP.E3 sgPair2), in comparison to the control, injected with Cas9 alone (zP.E3). The elastase-producing acinar cells of the exocrine tissue express mCherry (red) and nuclei are stained with DAPI (blue). Scale bar, 60 μm.

**Supplementary Figure 17.**
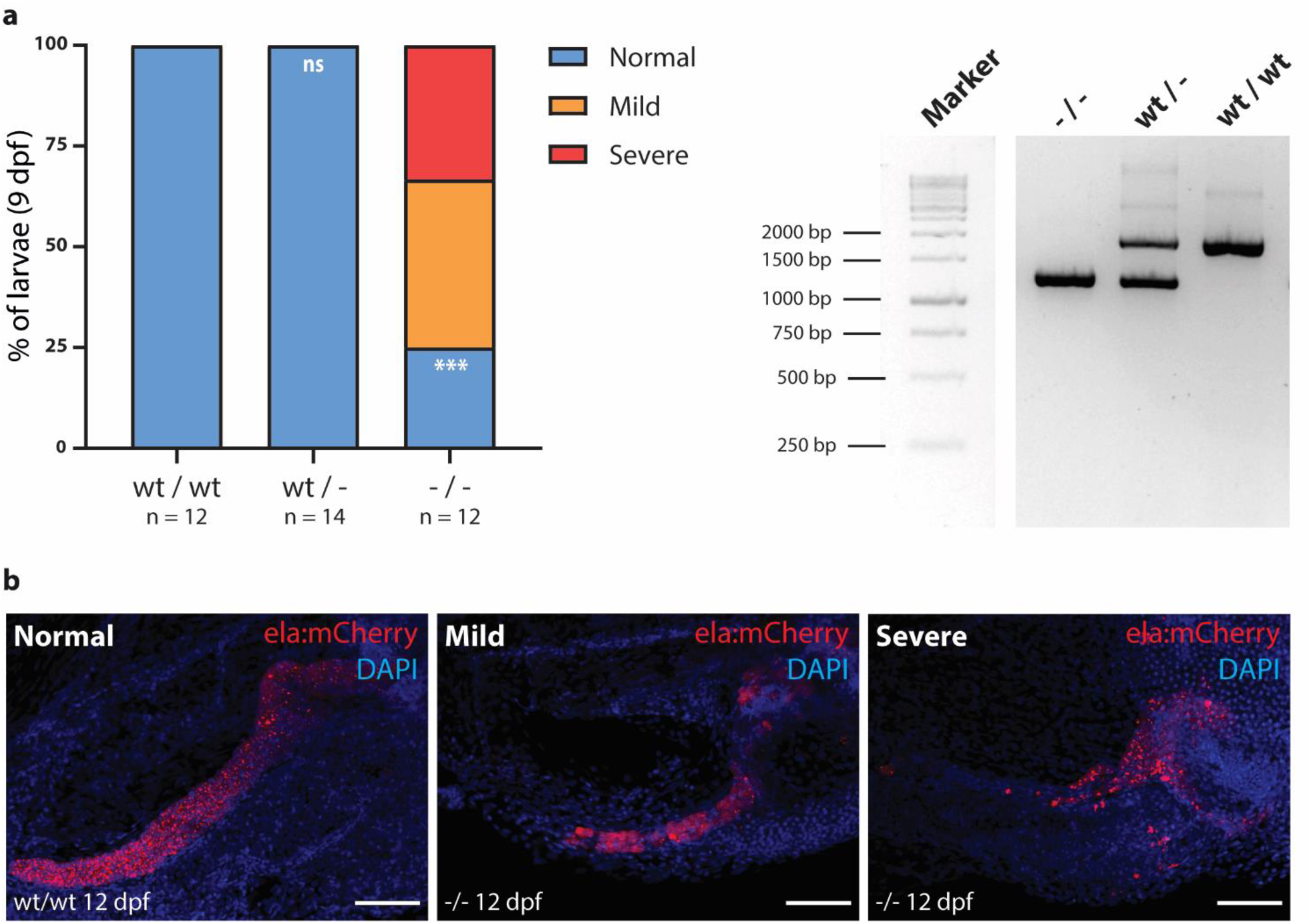
Homozygous deletion mutation in zP.E3 (zptf1aE3) affects pancreas size in F2 larvae. **a)** Homozygous embryos for the zP.E3 deletion (-/-) show mild to severe pancreatic defects, unlike wild type (wt/wt) and heterozygous (wt/-) sibling embryos (left). PCR screening of zPtf1aE3 deletion in F1 zebrafish embryos: wt/wt, wt/- and -/-. **b)** Representative confocal images (maximum intensity projections) of 12 dpf Tg(*elastase:mCherry*) larvae showing impaired development of pancreas upon CRISPR/Cas9-mediated deletions in zP.E3^-/-^ larvae compared to the control (zP.E3^wt/wt^). The elastase-producing acinar cells of the exocrine tissue express mCherry (red) and nuclei are stained with DAPI (blue). Scale bar, 60 μm.

